# Disentangling the Independent Contributions of Visual and Conceptual Features to the Spatiotemporal Dynamics of Scene Categorization

**DOI:** 10.1101/2020.04.11.037127

**Authors:** Michelle R. Greene, Bruce C. Hansen

**Affiliations:** Program in Neuroscience, Bates College; Department of Psychological & Brain Sciences, Neuroscience Program, Colgate University

## Abstract

Human scene categorization is characterized by its remarkable speed. While many visual and conceptual features have been linked to this ability, significant correlations exist between feature spaces, impeding our ability to determine their relative contributions to scene categorization. Here, we employed a whitening transformation to decorrelate a variety of visual and conceptual features and assess the time course of their unique contributions to scene categorization. Participants (both sexes) viewed 2,250 full-color scene images drawn from 30 different scene categories while having their brain activity measured through 256-channel EEG. We examined the variance explained at each electrode and time point of visual event-related potential (vERP) data from nine different whitened encoding models. These ranged from low-level features obtained from filter outputs to high-level conceptual features requiring human annotation. The amount of category information in the vERPs was assessed through multivariate decoding methods. Behavioral similarity measures were obtained in separate crowdsourced experiments. We found that all nine models together contributed 78% of the variance of human scene similarity assessments and was within the noise ceiling of the vERP data. Low-level models explained earlier vERP variability (88 ms post-image onset), while high-level models explained later variance (169 ms). Critically, only high-level models shared vERP variability with behavior. Taken together, these results suggest that scene categorization is primarily a high-level process, but reliant on previously extracted low-level features.

**Significance Statement:** In a single fixation, we glean enough information to describe a general scene category. Many types of features are associated with scene categories, ranging from low-level properties such as colors and contours, to high-level properties such as objects and attributes. Because these properties are correlated, it is difficult to understand each property’s unique contributions to scene categorization. This work uses a whitening transformation to remove the correlations between features and examines the extent to which each feature contributes to visual event-related potentials (vERPs) over time. We found that low-level visual features contributed first, but were not correlated with categorization behavior. High-level features followed 80 ms later, providing key insights into how the brain makes sense of a complex visual world.

## 1. Introduction

Human scene processing is characterized by its high speed: not only do observers require little viewing time to reliably understand scene content (M. R. Greene & Oliva, 2009; Potter, Wyble, Hagmann, & McCourt, 2014), but scene-specific neural responses have also been observed less than 200 ms after scene presentation (Bastin et al., 2013; Ramkumar, Hansen, Pannasch, & Loschky, 2016; Thorpe, Fize, & Marlot, 1996). However, we know comparatively little about the processing stages that transform the retinal image into a semantically rich categorical representation. Ongoing research has demonstrated that scene categories can be distinguished on the basis of many types of features, ranging from low-level visual properties such as histogram statistics of colors, edges, orientations, or Fourier metrics (Hansen & Loschky, 2013; Oliva & Schyns, 2000; Torralba & Oliva, 2003; Walther & Shen, 2014), to mid-level representations including texture (Renninger & Malik, 2004); “bag of words” representations describing the list of objects within scenes (Greene, 2013); or geometric properties of spatial layout (M. Greene & Oliva, 2009; Oliva & Torralba, 2001); to high-level properties such as conceptual attributes (Patterson, Xu, Su, & Hays, 2014) and affordances (Bonner & Epstein, 2018; Greene, Baldassano, Esteva, Beck, & Fei-Fei, 2016). However, we do not know the relative contributing strengths of each of these features to categorization, nor the time course of their contributions.

A powerful way to examine feature contributions is to consider each as a representational feature space (Edelman, 1998; Gärdenfors, 2004; Kriegeskorte et al., 2008). In this framework, each scene is considered a point in a high-dimensional space whose dimensions correspond to individual feature levels within the space. For example, in the feature space of *objects*, an office can be described by the presence of objects within it, such as “desk”, “monitor”, and “keyboard”. In the feature space of *texture*, the same scene would be described as a set of features describing the grain of wood on the desk, or the pattern of the carpet. Critically, such conceptual spaces can be used to make predictions about the types of errors that an observer or model will make about an image. For example, the *object* feature space would predict that images that share objects with offices would be frequently confused with offices (for example, a desk and monitor might be found in a college dorm room).

Despite the power of this approach, challenges remain in assessing the relative contributions of low- and high-level features (Groen, Silson, & Baker, 2017; Malcolm, Groen, & Baker, 2016), primarily because these features are not independent. Consider removing a stove from an image of a kitchen. This alteration not only changes the list of objects in the scene, but also changes the scene’s spatial layout as objects define the shape of a scene’s layout (Biederman, 1981). Furthermore, this change also alters the distribution of low-level visual features such as colors and orientations that belonged to the stove, and also changes the affordances of the space: it is much more difficult to cook without the stove. Altogether, these intrinsic correlations mean that we cannot easily interpret the use of any particular feature except in isolation from the others.

Here, we have addressed this problem by decorrelating a large number of predictive models that ranged from low-level visual properties to high-level semantic descriptors prior to analysis. Additionally, we leveraged an optimized category selection procedure that enabled maximal differentiation between the competing models across 30 different scene categories. Using high-density electroencephalography (EEG), we examined the relative power of each encoding model to explain the visual event-related potentials (vERPs) that are linked to scene categorization, as indexed via multivariate decoding and behavioral similarity assessments. Altogether, our results show a striking dissociation between feature processing and their use in behavior: while low-level features explain more overall vERP variability, only high-level features are related to behavioral responses.

## 2. Methods

### 2.1 Apparatus

All stimuli were presented on a 23.6” VIEWPixx/EEG scanning LED-backlight LCD monitor with 1 ms black-to-white pixel response time. Maximum luminance output of the display was 100 cd/m^2^, with a frame rate of 120 Hz and a resolution of 1920 x 1080 pixels. Single pixels subtended 0.0373 degrees of visual angle as viewed from 32 cm. Head position was maintained with a chin rest (Applied Science Laboratories).

### 2.2 Participants

We conducted the experiment twice, with a total of 29 observers volunteering across the two studies. Fourteen participants (6 female, 13 right handed) participated in the primary experiment. One participant’s EEG data contained fewer than half valid trials following artifact rejections and was therefore not included in subsequent analysis. Fifteen observers (9 female, 12 right handed) participated in an internal replication study (all results reported in Supplementary Materials and are largely consistent with the data presented here). The age of all participants ranged from 18 to 22 years (mean age = 19 years). All participants had normal or corrected to normal vision as determined by standard ETCRS acuity charts. The experimental protocol was approved by the Colgate University Institutional Review Board, and all participants provided written informed consent before participating, and were compensated for their time.

### 2.3 Stimuli

The stimulus set consisted of 2250 color photographs taken from 30 different scene categories (75 exemplars per category), within the SUN database (Xiao, Ehinger, Hays, Torralba, & Oliva, 2014). Category selection was conducted as to ensure maximally different representational dissimilarity matrices (RDMs) across three different feature types: visual features, defined as activations from the penultimate layer of a pre-trained deep convolutional neural network (Sermanet et al., 2013); object features, defined as a bag-of-words model over hand-labeled objects (Fei-Fei & Perona, 2005; Lazebnik, Schmid, & Ponce, 2006); and functional features, defined as hand-labeled scene affordances, taken from the American Time Use Survey (Greene et al., 2016). The optimization procedure was inspired by the odds algorithm of (Bruss, 2000). Specifically, we created 10,000 pseudorandom sets of 30 categories balanced across superordinate scene category (10 indoor, 10 urban outdoor, 10 natural landscape). RDMs for each of the three models were constructed, and the inter-model correlations were recorded. After this initial set of observations, we continued to create pseudo-random category sets until we observed a set with lower inter-model correlations than anything previously observed. The number of categories was determined by balancing the desire to represent the full diversity of visual environments, with the need to keep the experiment of manageable length.

We selected 75 images per each of the 30 scene categories. When possible, these were taken from the SUN database. In other cases, we sampled additional exemplars from the internet (copyright-free images). Care was taken to omit images with salient faces in them. All images had a resolution of 512 x 512 pixels (subtending 20.8 degrees of visual angle) and were processed to possess the same root-mean-square (RMS) contrast (luminance and color) as well as mean luminance. All images were fit with a circular linear edge-blurred window to obscure the square frame of the images, thereby distributing contrast changes around the circular edge of the image (Hansen & Essock, 2004).

### 2.4 Human Scene Category Distance Measurement

In order to model category distances from human behavior, we conducted a series of six experiments on Amazon’s Mechanical Turk marketplace. This was necessary because the long image presentation time in the EEG experiment (750 ms) led to ceiling-level categorization performance. Each behavioral experiment assessed observers’ judgments of scene similarity by presenting three items and asking the observer to choose the odd-one-out. Although this task specifically queries similarity, it has recently been shown to reveal hierarchical category representations of objects (Zheng, Pereira, Baker, & Hebart, 2019). Thus, we use it here as a measure of scene categorization behavior.

The first experiment queried 608 participants about scene similarity without constraining the definition of similarity. The other five experiments asked participants to determine scene similarity with respect to one of five features: global orientation (N=202), texture (N=176), objects (N=104), functions/affordances (N=99), and lexical (N=820).

For all experiments, participants were selected from a pool of United States-based workers who had previously completed at least 1000 hits with an approval rating of at least 95%. Each participant was able to complete as many hits as they wished, and each hit consisted of 20 trials. Each participant completed between 1 and 227 hits (median: 2 hits). We collected a total of 5000 hits for the unconstrained similarity experiment, and 1000 in each of the other five experiments. Thus, a minimum of 24 participants rated each category triad in the unconstrained experiment, and 5 participants per triad in the remaining experiments.

With the exception of the lexical similarity experiment, all experiments were identical though each had a different definition of similarity. Each trial consisted of three images from three unique categories. These images were presented side by side in a single row. The participant was instructed to click on the image that was the least similar to the other two, given the particular similarity instructions for that experiment (Zheng et al., 2019). For the lexical experiment, images were replaced with the name of the scene category on a blank gray background. Each hit was completed in a median work time ranging from 104 seconds in the texture experiment to 159 seconds in the orientation experiment, and participants were compensated $0.10 per hit for their time.

From these responses, for each experiment, we created a 30-category by 30-category distance matrix as follows. Beginning with a 30 x 30 matrix of zeros, for each trial of each hit, we added one to the matrix entry representing the row of the selected category and each column of the two alternatives. This represents the participant’s judgment that the selected image was dissimilar to the other two. We then subtracted one from the matrix entry representing the non-selected alternatives. This represents the participant’s judgment that the two non-selected images were deemed to be more similar to one another. The final distance matrix was normalized to be between 0 and 1, and the off-diagonal entries were saved to become a regressor in subsequent analyses.

### 2.5 EEG Experimental Procedure

Participants performed a three-alternative forced choice (3AFC) categorization task on each of the 2250 images across two ~50-minute recording sessions. All images within each category were randomly split into two sets, and the image set was counterbalanced across participants. Within both sets, image order was randomized.

Each trial commenced with a 500 ms fixation followed by a variable duration (500-750 ms) blank mean luminance screen to allow any fixation-driven activity to dissipate. Next, a scene image was presented for 750 ms followed by a variable 100-250 ms blank mean luminance screen, followed by a response screen consisting of the image’s category name and the names of two randomly selected distractor category names presented laterally in random order. Observers indicated category choice by clicking on the appropriate category name with a mouse.

### 2.6 EEG Recording and Processing

High-density 256-channel EEGs were recorded in a Faraday chamber using Electrical Geodesics Incorporated’s (EGI; Phillips Neuro) Geodesic EEG acquisition system (GES 400) with Geodesic Hydrocel sensor nets (electrolytic sponges). The online reference was at the vertex (Cz), and the impedances were maintained below 50*k*Ω (EGI amplifiers are high-impedance). All EEG signals were amplified and sampled at 1000 Hz. The digitized EEG waveforms were first highpass filtered at a 0.1 Hz cut-off frequency to remove the DC offset, and then lowpass filtered at a 45 Hz cutoff frequency to eliminate 60 Hz line noise.

All continuous EEGs were divided into 850 ms epochs (100 ms before stimulus onset and 750 ms of stimulus-driven response). Trials that contained eye movements or eye blinks during data epochs were excluded from analysis. Additionally, all epochs were subjected to algorithmic artifact rejection whereby voltages exceeding +/- 100*μV* or transients greater than +/- 100*μV* were omitted from further analysis. These trial rejection routines resulted in no more than 10% of trials being rejected from any one participant. Each epoch was then re-referenced offline to the net average, and baseline-corrected to the last 100 ms of the blank interval that preceded the image interval. Grand average vERPs were assembled by averaging all re-referenced and baseline-corrected epochs across scene category and participants.

For both encoding and decoding analyses, we improved the signal-to-noise ratio of the single trial data by building ‘supertrials’ by averaging 20% of trials within a given category, drawn randomly without replacement (e.g. (Cichy, Khosla, Pantazis, Torralba, & Oliva, 2016; Isik, Meyers, Leibo, & Poggio, 2014)). This process was repeated separately for each participant. This approach is desirable as we are primarily interested in category-level neuroelectric signals that are time-locked to the stimulus.

Topographic plots were generated for all experimental conditions using EEGLAB (Delorme & Makeig, 2004) version 13.5.4b in MATLAB (version 2016a, The Mathworks, Natick, MA). Source localization was conducted via EGI’s GeoSource 3.0 source-imaging package and corresponding Geodesic Photogrammetry System (GPS). Individual head models were obtained from each participant using the photogrammetry system and solved using GeoSource 3.0 software. The dense array of Geodesic sensor locations obtained from each participant enables high-resolution finite difference method (FDM) conformal MRI atlas head models (Li, Papademetris, & Tucker, 2016), and demonstrably high source localization accuracy (Kuo et al., 2014; Song et al., 2015). Here, the inverse problem was solved using the inverse mapping constraint LORETA with a regularization *α* = 3.

### 2.7 Decoding Procedure

Prior to analysis, all data were downsampled to 500 Hz to speed computation. For each participant, scene category decoding was conducted on an electrode-by-electrode basis in 41 ms windows centered at each time point (i.e. +/- 20 ms around and including the given time point). Given that the window could not extend beyond the 750 ms image period, the analysis was truncated at 730 ms. Scene category decoding was conducted using a linear one-versus-all multi-class support vector machine (SVM), implemented in Matlab’s Statistics and Machine Learning Toolbox (Version 10.0). Accuracy of the SVM decoder was measured using 5-fold cross-validation, and empirical 95% confidence intervals for decoding accuracy were calculated across participants along the main diagonal of the 30 x 30 decoder confusion matrix.

### 2.8 Encoding Models

We employed a total of 9 different encoding models, representing a range of visual and conceptual features. Models were chosen for inclusion in this study because they have been shown to explain significant variance in brain or behavioral data, rather than for biological plausibility per se. The models fall into four broad classes: three models represent outputs of multi-scale Gabor wavelet pyramids, or their derivatives (Wavelet, Gist, and Texture); two models representing activation outputs from a deep convolutional neural network (dCNN) that used the eight-layer “AlexNet” architecture (Krizhevsky, Sutskever, & Hinton, 2012) and was pre-trained to perform scene classification on the Places database (Zhou, Lapedriza, Khosla, Oliva, & Torralba, 2017). For these models, we chose one lower-level convolutional layer (Conv2), as well as one higher-level fully-connected layer (FC6). In order to represent higher-level visual information that we cannot yet obtain directly from images, three models were included whose features were obtained by human ratings (Objects, Attributes, and Functions). Last, we considered the semantic similarity across scene categories, operationalized as the lexical distance between category names. For each model except lexical distance, representational dissimilarity matrices (RDMs) were created by computing the distance between each pair of categories in the given feature space using the 1 - correlation distance metric (Spearman’s *ρ*).

#### 2.8.1 Filter Models

##### 2.8.1.1 Wavelets

In order to encode the low-level structural details of each scene, we used the outputs of a multi-scale bank of log-Gabor filters (Field, 1987) that decompose an image by spatial frequency, orientation, and spatial location. Such encoders are well-established models of early visual cortex (e.g. (Carandini et al., 2005)), as well as front-ends to machine vision systems (Simoncelli & Freeman, 1995). Each image was converted from RGB to L*a*b color and passed through a bank of log-Gabor filters at seven spatial frequencies (0.125, 0.25, 0.5, 1, 2, 4, and 8 cycles per degree) and four orientations (0, 45, 90, and 135 degrees). The filters had a spatial frequency bandwidth of approximately 1.5 octaves, assessed at full width at half height. Thus, each of the three L*a*b channels was represented by 28 filter outputs, for a total of 84 features per image. We averaged features across images within a category to create a 30-category by 84-feature matrix.

##### 2.8.1.2 Gist

Each image was described with the spatial envelope (or Gist) descriptor of (Oliva & Torralba, 2001). These features represent a summary statistic representation of the dominant orientation contrast at three different spatial frequencies localized in a 8 x 8 window grid across the image. The number of orientations varies with spatial frequency, with 8 orientations at the highest frequency, 6 in the mid-range, and 4 at the lowest, for a total of 1152 features (64 windows x (8+6+4) orientations per scale). These features have been shown to explain significant variance in fMRI and MEG responses throughout visual cortex (Ramkumar et al., 2016; Watson, Andrews, & Hartley, 2017; Watson, Hartley, & Andrews, 2014). Given their similarity to the log-Gabor filters described above, following (Oliva & Torralba, 2001), we used linear discriminant analysis (LDA) to learn weights on the filter outputs that were optimized for classifying the 30 scene categories of this database. We averaged across images in each category to create a 30-category by 1152-feature matrix.

##### 2.8.1.3 Texture

Texture features for each image were encoded using the generative texture model of (Portilla & Simoncelli, 2000). This algorithm analyzes a total of 6495 statistics from an image, including marginal pixel statistics, wavelet coefficient correlations, wavelet magnitude correlations, and cross-scale phase statistics. Scenes can be distinguished by their texture properties (Renninger & Malik, 2004), and texture properties have been reported to drive activity in both early visual areas such as V2 (Freeman & Simoncelli, 2011), as well as the parahippocampal place area (PPA, (Cant & Goodale, 2011; Lowe, Rajsic, Gallivan, Ferber, & Cant, 2017)). As before, we averaged across images in a category to create a 30-category by 6495-feature matrix.

#### 2.8.2 Deep CNN Features

We extracted the activations from two of the eight layers in a deep convolutional neural network (Conv2 and FC6 from a CNN trained on the Places database (Zhou et al., 2017), using the AlexNet architecture (Krizhevsky et al., 2012), and implemented in Caffe (Jia et al., 2014). This CNN was chosen because it is optimized for classification of 205 scene categories and because this eight-layer architecture is most frequently used when comparing CNNs to brain activity (Bonner & Epstein, 2018; Cadieu et al., 2014; Cichy, Khosla, Pantazis, & Oliva, 2017; Cichy et al., 2016; Greene & Hansen, 2018; Güçlü & Gerven, 2015; Khaligh-Razavi & Kriegeskorte, 2014; Kubilius, Bracci, & Beeck, 2016). We chose Conv2 and FC6 as the layers of interest because they have been argued to represent typical low- and high-level feature information respectively and have previously demonstrated fundamentally different encoding behavior with EEG data (Greene & Hansen, 2018). In the Conv2 layer, 256 5 x 5 pixel filters are applied to the input with a stride of one pixel, and padding of two pixels, for a total of 27×27×256 = 186,624 features. The sixth and seventh layers of AlexNet are fully-connected, meaning that they have no retinotopic information, and were designed to have 4096 features each. For both layers, we averaged across images within each category, creating a 30-category by 186,624-feature matrix for Conv2 and a 30-category by 4096-feature matrix for FC6.

#### 2.8.3 Models of Conceptual Features

##### 2.8.3.1 Objects

We employed a bag-of-words object representation used in (Greene et al., 2016). Each object or region in each scene was hand-labeled using the LabelMe tool (Russell, Torralba, Murphy, & Freeman, 2008). Objects are features that can be diagnostic of a scene category (Greene, 2013), and have been reported to drive activity across occipitotemporal cortex (Harel, Kravitz, & Baker, 2012; MacEvoy & Epstein, 2011). Across the image set, there were 3563 unique object labels. We created a 30-category by 3563-object matrix that stores the proportion of scenes in each category *i* containing each object *j*.

##### 2.8.3.2 Scene Attributes

We took the measured attribute vectors of (Patterson et al., 2014). The attributes were obtained by asking human observers to list features that made pairs of images different from one another in a massive online norming experiment. The resulting attributes consist of a heterogeneous set of 112 descriptors including aspects of region, material, and spatial layout. In a second round of normative ratings, Patterson and colleagues asked human observers on Amazon’s Mechanical Turk (AMT) to rate each scene in the SUN database according to 112 different attributes. Thus, each scene’s attribute rating is the proportion of mTurk observers selecting that attribute as appropriate for that scene. As before, we averaged attribute descriptions across category, resulting in a 30-category by 112-feature matrix.

##### 2.8.3.3 Functions/Affordances

Scene functions were operationalized as a description of the set of actions that a person could perform in each environment. As with the attributes, these vectors were also created by workers on AMT who annotated which of 227 actions in the American Time Use Survey could be done in each of the scenes. Scene functions have been shown to explain the majority of variance in scene categorization behavior (Greene et al., 2016; Groen et al., 2018). The final feature matrix consisted of a 30-category by 227-feature matrix in which each cell stored the proportion of images in a given category *i* were linked to a specific action *j*.

#### 2.8.4 Model of Semantics

##### Lexical Distance

Last, we included a model of semantic distance between category names, as semantic distance has been demonstrated to affect early processing and reaction times to objects and scenes (Neely, 1977). The semantic distance between categories was operationalized as the shortest path between category names in WordNet (Miller, 1995), as implemented by the Wordnet::Similarity tool (Pedersen, Patwardhan, & Michelizzi, 2004).

### 2.9 Time-Resolved Encoding Procedure

In order to compare vERP trial data to each of the models in a common framework, we employed representational dissimilarity analysis (RSA, (Kriegeskorte et al., 2008)). With this approach, we examined category similarity with respect to each of the nine feature spaces to scene category similarity with respect to time-resolved vERP activity at each electrode.

#### 2.9.1 Model Representational Dissimilarity Matrices (RDMs)

For each of the nine models, we created a 30 x 30-category correlation matrix from the feature matrices described above. Representational dissimilarity matrices (RDMs) were defined as 1 - Spearman’s *ρ*. Importantly, RDMs are symmetric with an undefined diagonal. Therefore, only the lower triangle of each RDM was included in the analysis, representing 435 pairs of category distances. As shown in **Figure 1**, substantial correlations exist between each of the nine models. Therefore, we used a whitening transformation (Bell & Sejnowski, 1997) in order to decorrelate the feature spaces. Specifically, for each feature vector x, we sought to obtain a new feature vector z that is uncorrelated to the other feature vectors. The linear transformation that can achieve this is:

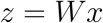

**Figure 1:**
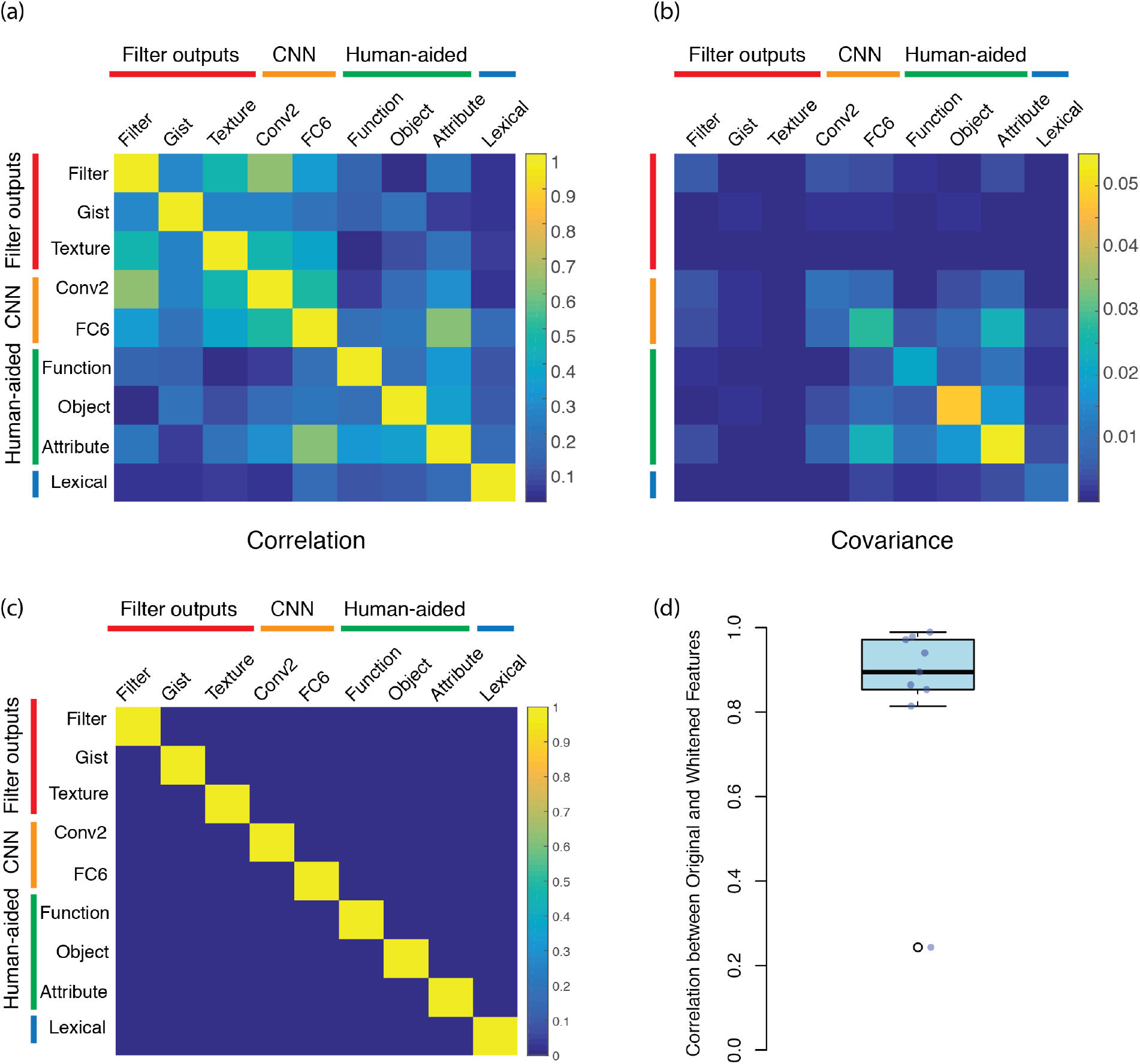
(a) Correlations between the nine original features; (b) Covariance before ZCA whitening transformation; (c) Correlations after ZCA whitening transformation; (d) Correlations between original and whitened features. Gist features are the significant outlier.

In order to decorrelate each feature, the whitening matrix *W* must satisfy:

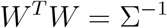

Under these conditions, the covariance matrix Σ of z is equal to the identity matrix. However, there are multiple whitening matrices that would achieve this transformation. Following (Kessy, Lewin, & Strimmer, 2017), we opted for the zero-phase component algorithm (ZCA) as it was found to provide sphered components that were maximally similar to the original data. In the ZCA algorithm, the whitening matrix is forced to be symmetrical (i.e. *W^T^* = *W*), and *W* = Σ^−1/2^. In order to visualize the scene category similarity structure for each of the nine whitened feature spaces, **Figure 2** shows two-dimensional multidimensional scaling (MDS) solutions for each feature space.

**Figure 2:**
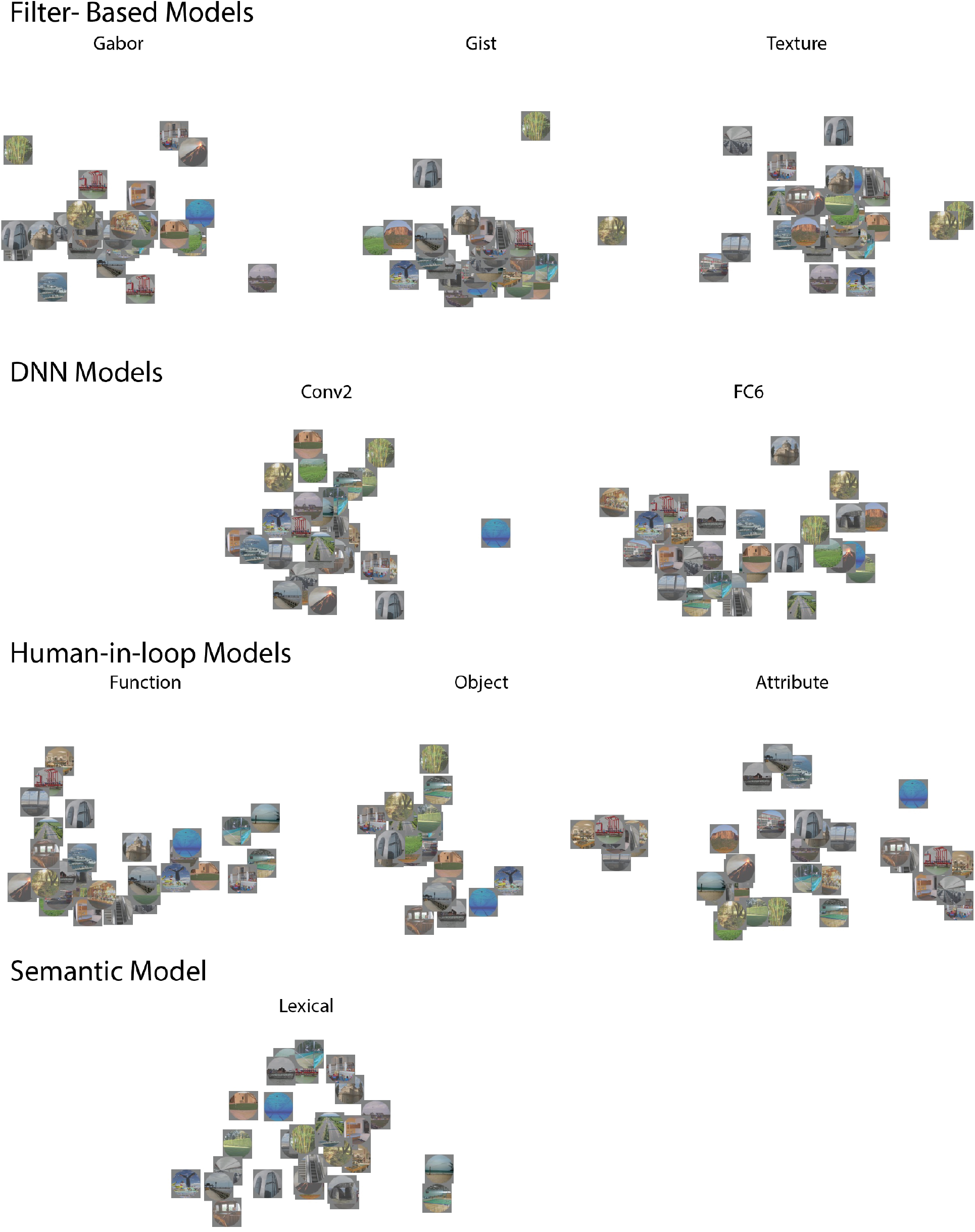
Representation of each of the nine whitened feature spaces. To aid visualization, one representative image from each of the 30 categories is used to plot the category’s location in a 2D multidimensional scaling (MDS) solution.

#### 2.9.2 Neural RDMs

For each participant, we averaged vERPs across trials within the same category, and then normalized the resulting averages. For each electrode, we extracted vERP signals within a 41 ms window centered on each time point, beginning 100 ms before scene presentation, and extending to the entire 750 ms of stimulus presentation, truncating the window as necessary if the entire 41 ms was not available in an epoch. Thus, each window consisted of a 41 time-point by 30 category matrix. From this matrix, we created a 30 x 30 RDM using the same 1 - Spearman metric described above. As before, the lower triangle of this matrix (435 points) was used as the dependent variable in the regression analyses.

#### 2.9.3 Computing Noise Ceiling

In order to quantitatively assess model fits, we computed the noise ceiling of our data, following the methods of (Nili et al., 2014). Briefly, the upper bound of the noise ceiling is an overfit value representing the explained variability of the group mean to predicting an individual observer whose data are included in the group mean. The lower bound represents the explained variability of the group mean when the predicted observer is left out of the group mean. This procedure was performed independently at each time point.

### 2.10 Electrode Selection

As the primary goal of this study is to examine the genesis of scene category representations, for the encoding analyses, we selected only the electrodes containing significant category information from decoding. Specifically, we used the decoding accuracies for each electrode to select electrodes for encoding analyses that had accuracy values above the 95% confidence interval of the pre-stimulus baseline for at least 10 continuous milliseconds. This amounted to an average of 248 electrodes per participant (range: 232-256). We averaged across these electrodes for all subsequent encoding analyses.

### 2.11 Statistical Analysis

For all encoding analyses, we used a jackknife approach to obtain stable maximum R^2^ and latency of maximum R^2^ values for each participant. This was done by iteratively leaving out one of the participants in turn, and then computing the statistic of interest. The maximum R^2^ values for each participant were therefore defined as the maximum R2 from the mean of the remaining 12/13 participants, and the latency of the maximum was defined as the time point when this maximum was observed. All F and t values were corrected using the methods suggested by (Luck, 2005): namely, that *t* values were divided by *N* – 1 and F values were divided by (*N* – 1)^2^. Group-level significance was assessed via sign test. Post-hoc t-tests were corrected for multiple comparisons across time points and models using the Benjamini-Hochberg procedure.

## 3. Results

We begin by establishing the role of our nine whitened features in human scene categorization (Section 3.1) and show the consistency of our vERP results with the existing literature (Section 3.2). We then establish the availability of scene category information in vERP data using time-resolved decoding (Section 3.3). Finally, we assess the extent to which the nine whitened features map to vERP data over time and across task (Section 3.4 and beyond) in order to gain insight into the representational transformations that enable rapid scene categorization.

### 3.1 Feature Use in Scene Categorization Behavior

#### 3.1.1 Predicting Unconstrained Similarity Judgment from Feature Spaces

We began by establishing the extent to which observers utilize each of the nine whitened feature models in the unconstrained similarity task. As the long stimulus presentation time (750 ms) in the EEG experiment led to ceiling-level categorization performance, we used the scene similarity results from mTurk (see Section 2.4) as a proxy for categorization behavior. Although each of the nine feature models in this study have been strongly associated with scene categorization behavior, this first step ensures that this holds when the features have been whitened. Further, as most studies only examine a few features in isolation, this analysis shows the extent to which each feature contributes to scene category representations.

Collectively, the nine whitened feature RDMs predicted 78% of the variance (adjusted R^2^) in the behavioral RDM obtained from the Mechanical Turk participants in the unconstrained experiment (F(9,425) = 163.8, p<0.0001). The beta coefficients, partial R^2^, and *p* values for each of the RDMs are shown in **Table 1**.

**Table 1:** Regression coefficients, partial R^2^, and p values for each of nine feature RDMs in predicting dissimilarity matrix from human observers’ rankings.

| Feature | Beta | Partial.R2 | p |
| --- | --- | --- | --- |
| Wavelets | 0.014 | 0.0160 | 0.009 |
| Gist | 0.005 | 0.0020 | 0.34 |
| Texture | 0.011 | 0.0090 | 0.047 |
| Conv2 | 0.003 | 0.0008 | 0.55 |
| FC6 | 0.121 | 0.5430 | <2e-16 |
| Functions | 0.080 | 0.3390 | <2e-16 |
| Objects | 0.034 | 0.0840 | 9.70E-10 |
| Attributes | 0.141 | 0.6150 | <2e-16 |
| Lexical | 0.024 | 0.0440 | 1.09E-05 |

Most whitened feature RDMs had significant predictive power for the unconstrained behavioral RDM. The two exceptions were the Gist features and the early dCNN layer (Conv2). However, as the Gist features were most transformed by the whitening procedure (see **Figure 1**), we are not strongly interpreting this result. Lower-level features, including the filter-based models and the early dCNN model were less predictive of unconstrained similarity judgments than the higher-level models. Replicating previous observations, the function-based model was more predictive of scene similarity assessments than the object-based model (Greene et al., 2016; Groen et al., 2018). However, the two most predictive models were the upper-layer features of the dCNN (FC6) and the attributes model of (Patterson et al., 2014).

The attribute model is itself a combination of four different feature types: affordances, materials, surfaces, and spatial properties. Therefore, it is not terribly surprising that it did so well when compared to individual feature spaces. In order to break down its predictive power, we created four separate RDMs corresponding to the four different types of attributes outlined by (Patterson et al., 2014): functions/affordances (36 features), materials/objects (36 features), surface properties (12 features), and spatial layout properties (14 features). We used these four as regressors for the human response RDM as before. Collectively, the four aspects of the attributes predicted 68% of the variance (adjusted R2) in the human RDM (F(4,430) = 231.3, p<<0.0001). **Table 2** shows beta coefficients, partial R2 and p values for each of the four.

**Table 2:** Regression coefficients, partial R^2^, and p values for each of the four attribute types that constituted the full attributes model.

| Model | Beta | Partial.R2 | p |
| --- | --- | --- | --- |
| Affordances | 0.430 | 0.35 | <2e-16 |
| Materials | 0.170 | 0.07 | 1.22E-08 |
| Surfaces | 0.054 | 0.02 | 0.00559 |
| Spatial Envelope | 0.170 | 0.12 | 4.28E-14 |

Overall, affordances were the most predictive attribute type, followed by the “spatial envelope” properties which consisted of spatial layout properties such as *openness*, *mean depth*, and level of *clutter*. By contrast, material properties such as *vegetation, asphalt*, or *metal*, as well as surface properties such as *dry, aged*, or *dirty*, contributed comparatively little to the behavioral RDM. Altogether, these results validate the use of these nine models for explaining variability in scene similarity and categorization behavior, consistent with previous results.

#### 3.1.2 Predicting Feature-Based Similarity Judgments from Feature Spaces

Next, we examined how changing the observer’s similarity task alters feature use in the five feature-based similarity experiments. These similarity experiments yielded RDMs that were highly correlated with one another (mean r=0.82, range: 0.67-0.94). In order to focus on the independent predictions made by these experiments, we combined the RDMs for each of the five experiments into a single data matrix (435 x 5) and performed the same whitening transform on this matrix, using the same procedure that we employed for the feature matrix (see Section 2.9.1). The resulting whitened RDMs were all highly correlated with the original results (mean r=0.73, range: 0.66-0.85), while becoming uncorrelated with one another.

As with the unconstrained similarity task, each of the five task-driven experiments were well-predicted by the nine whitened features: R^2^ values ranged from 0.61 for the lexical task to 0.75 for both functions/affordances and object tasks. **Table 3** shows partial R^2^ for each feature for each of the five task-driven experiments. In general, we observed that in each experiment, the pattern of feature use was highly correlated with the unconstrained similarity task (range: r=0.95 for orientation and lexical to r=0.99 for objects and functions). That said, there were also substantial task-driven differences, with higher partial R^2^ values observed for features that were task-relevant (wavelets for the orientation task, functions for the function/affordance task, and lexical features in the lexical task). One exception seems to be texture and object that appear to be switched in importance.

**Table 3:** Partial R^2^ for each feature (row) in each task-driven experiment (column).

| Feature | Orientation | Texture | Object | Functions | Lexical |
| --- | --- | --- | --- | --- | --- |
| Wavelet | 0.0570 | 0.01300 | 0.013 | 0.0170 | 0.00700 |
| Gist | 0.0120 | 0.00001 | 0.002 | 0.0002 | 0.00400 |
| Texture | 0.0008 | 0.00090 | 0.019 | 0.0030 | 0.00002 |
| Conv2 | 0.0170 | 0.02600 | 0.006 | 0.0280 | 0.01900 |
| FC6 | 0.5260 | 0.49800 | 0.542 | 0.5260 | 0.41500 |
| Functions | 0.1210 | 0.14700 | 0.251 | 0.2620 | 0.20300 |
| Objects | 0.0580 | 0.10800 | 0.009 | 0.0900 | 0.04000 |
| Attributes | 0.4660 | 0.59700 | 0.566 | 0.5690 | 0.29800 |
| Lexical | 0.0170 | 0.00300 | 0.040 | 0.0240 | 0.10700 |
| Total R2 | 0.7000 | 0.74000 | 0.750 | 0.7500 | 0.61000 |

In order to understand feature use across the five experiments, we isolated the five most relevant features from the nine feature spaces: wavelets, which should be most similar to the *orientation* task, texture statistics for the *texture* task, object features for the *object* task, functions for the *functions / affordances* task, and lexical for the *lexical* task. Figure 3 shows the regression coefficients for each of the five features (different plots) across the five experiments (different bars). With the exception of the object features, each feature had the highest regression weight for the predicted experiment. This result establishes that feature use differed across experimental task, and that specifically, a feature was used more when observers were asked to assess scene similarity with respect to that feature.

**Figure 3:**
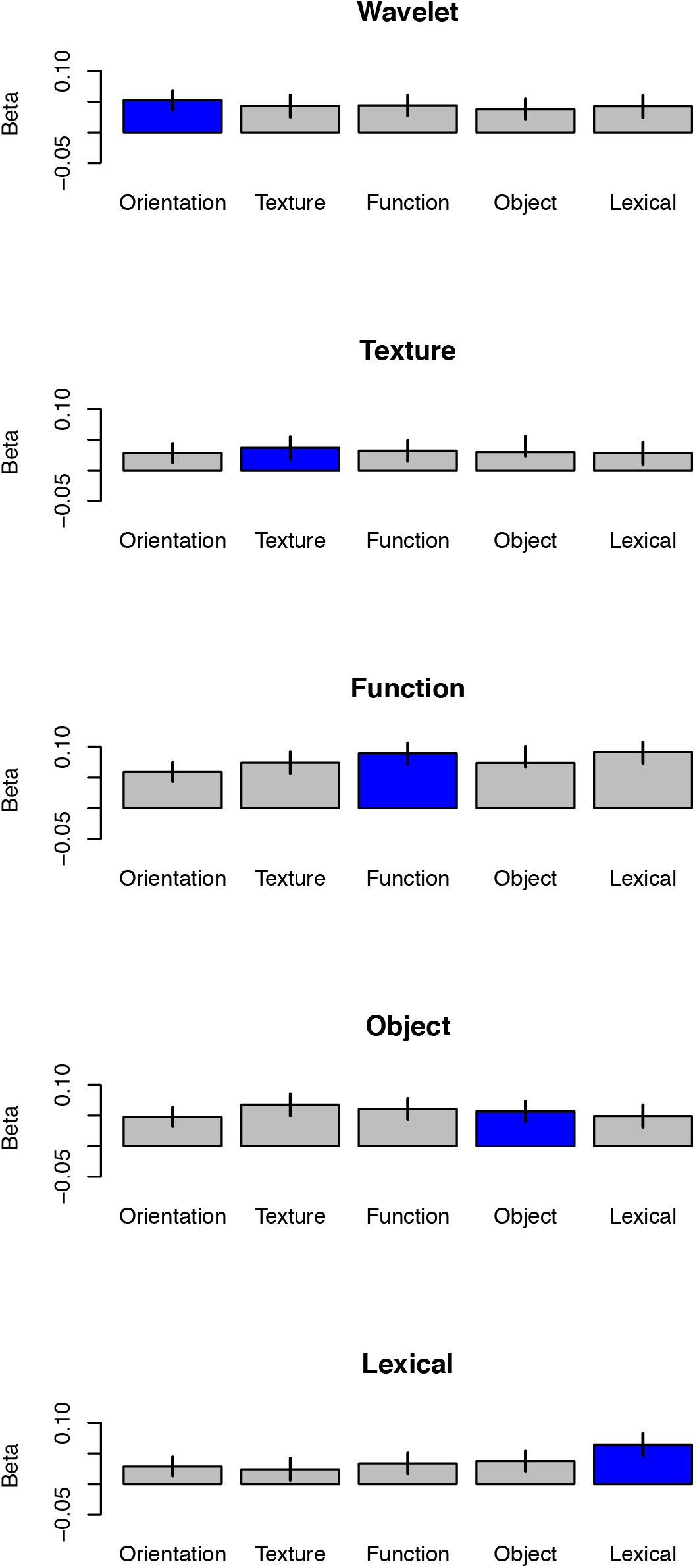
Feature use across the five similarity experiments. Each plot is a different feature, and the regression weight for that feature is shown across each of the five experiments. The blue bar indicates the experiment with the highest predicted weight. Error bars represent 95% confidence intervals.

### 3.2 vERP Data Summary

In this section, we examined the general spatiotemporal structure of the vERPs. Grand average vERPs and topographic plots are shown in **Figure 4a**. The topographic plots show the typical voltage difference scalp topography for observers engaged in viewing complex visual scenes (e.g. (Groen, Ghebreab, Lamme, & Scholte, 2012; Hansen, Jacques, Johnson, & Ellemberg, 2011; Hansen, Johnson, & Ellemberg, 2012)). **Figure 4b** shows the source localization results at select time points. The spatiotemporal evolution of the vERP sources is consistent with previous MEG studies in scene perception (e.g. (Ramkumar et al., 2016)), and show a gradual progression over time from primary visual cortices through bilateral occipito-temporal cortices, with apparent anterior temporal and ventral frontal cortices late. Therefore, the similarity between the spatiotemporal features of the current data with previously reported results contextualizes the subsequent modeling and decoding results within the broader M/EEG literature.

**Figure 4:**
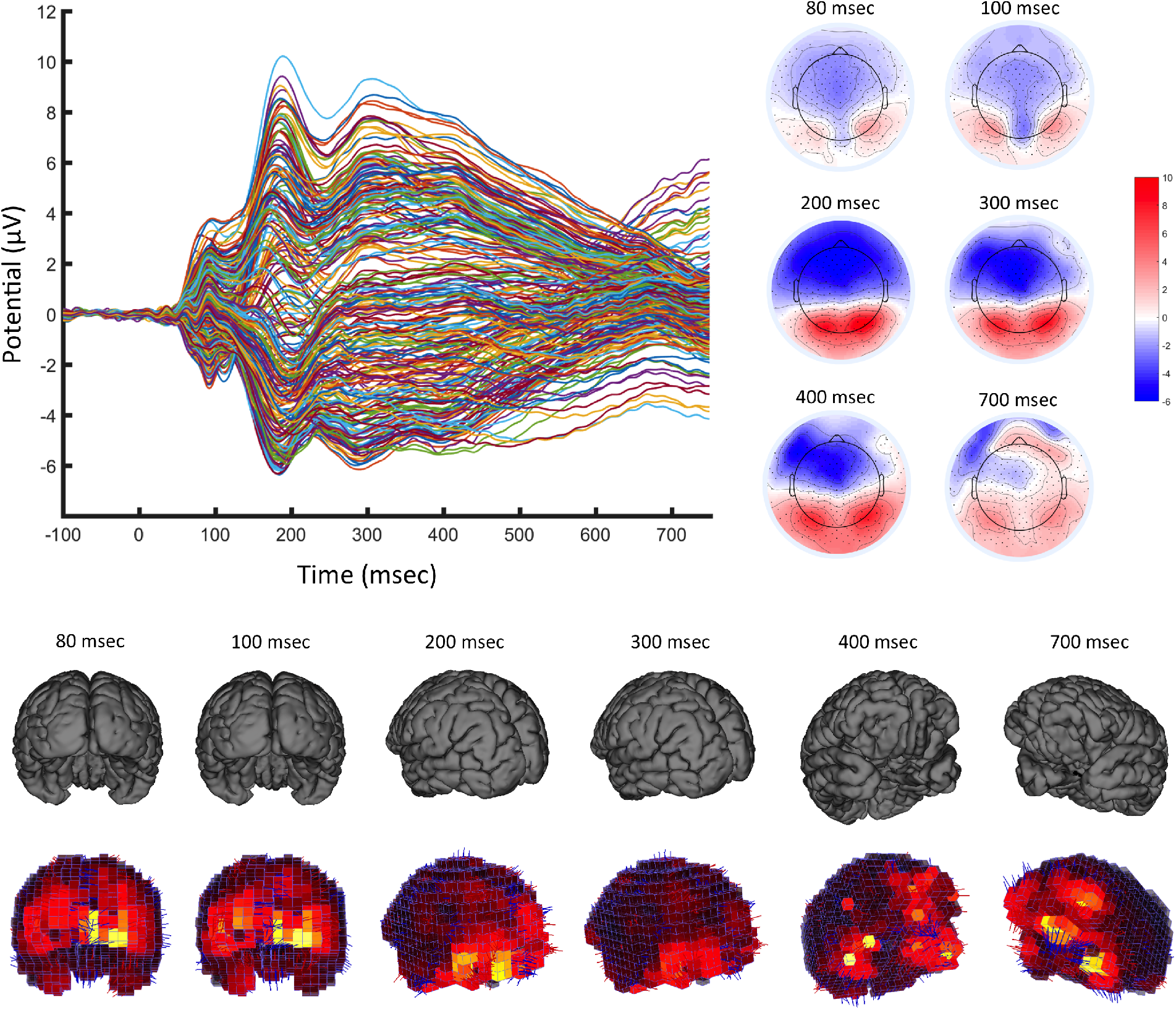
(a) Grand averaged vERPs (left), and topographic plots (right) for key time points. (b) Source-localization solutions from times ranging from 80 to 700 ms post-stimulus.

### 3.3 Time-Resolved Decoding

Our main goal is to examine the representational transformations that take place in the visual system en route to categorization. In order to measure the amount of scene category information available in vERPs over time, we employed a time-resolved decoding procedure on vERPs. **Figure 5** (and **Figures 5-1** and **5-2** for the replication experiment) show decoding performance across all electrodes over time. Significant category decoding was observed in nearly all electrodes, with an average of 248 of the 256 electrodes showing significant category information. Category information was highest between 100 and 200 ms after stimulus onset, with the maximum decodable information found on average 198 ms after stimulus onset (range: 142-214 ms). This corroborates previous accounts of generalized categorization taking place at or around 200 ms post-stimulus onset.

**Figure 5:**
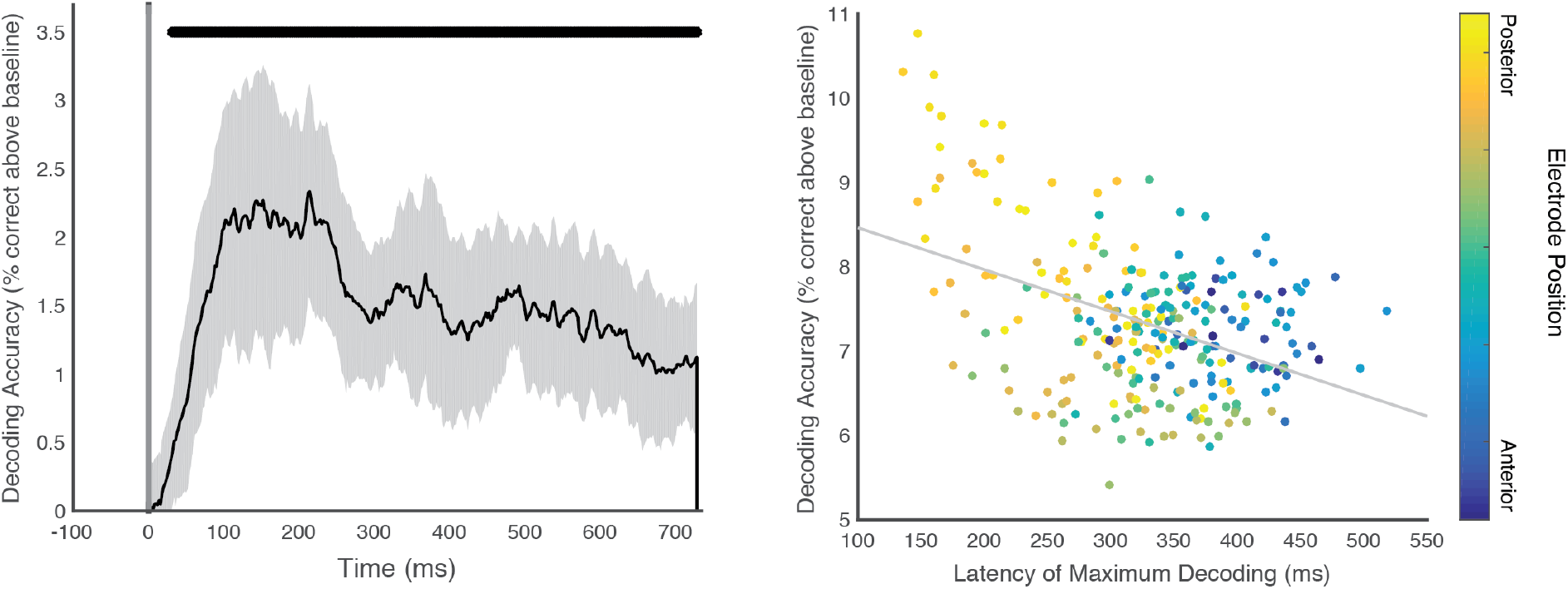
Left: Decoding performance relative to the false-positive rate observed during the baseline windows. Top line indicates decoding performance that is significantly over the false positive rate. Shaded area indicates 95% confidence interval. Right: Latency of maximum decoding performance and maximum decoding accuracy relative to the false-positive rate. Each point is an electrode, averaged across participants. Color represents cap location of electrode from posterior to anterior. Gray line represents the regression line.

Decoding analyses were performed on each electrode individually, allowing us to examine the pattern of decodable information across the scalp. Specifically, we examined the differences in decoding across electrodes by computing: 1) the maximum decoding accuracy at each electrode; and 2) the temporal latency of this maximum value. In addition, we ordered electrode position from anterior to posterior. We observed a sizable negative correlation (r = −0.42, 95% CI: −0.52 to −0.31, t(12) = 2.71, p < 0.05, see **Figure 5**) between the maximum decoding performance and the latency of maximum decoding for an electrode. This was also observed in the replication experiment (r=-0.42, p<0.0001, see **Figure 5-2**). This may suggest that electrodes that carry the most category information also have this information available earlier, or that later neural responses are less time-locked to stimulus presentation, driving down the amount of decodable category information. Further, we found that the overall decoding performance was correlated with electrode location from anterior to posterior (r = 0.21, 95% CI: 0.08 to 0.32, t(12) = 4.0, p<0.001, see **Figure 5**), indicating that decodable category information was concentrated in posterior electrodes. An even more pronounced effect was observed in the replication experiment (r=0,45, p<0.0001, see **Figure 5-2**).

### 3.4 Time-Resolved Encoding

#### 3.4.1 All Features

While the decoding analyses establish the amount of specific scene category information available in the vERPs, the goal of the encoding analyses is to establish the extent to which a variety of visual and conceptual features collectively explain variability in the vERPs. Taken together, the nine whitened models predicted significant vERP variability in electrodes with significant category information (see **Figure 6**). The maximal explained variability occurred 98 ms after stimulus onset, on average, and was within the noise ceiling for nearly the entire epoch. Interestingly, the explained variance of the features was below the noise ceiling between 176 and 217 ms post-image onset, the same time window of maximum category decoding. This may suggest that while these nine features are explaining the perceptual processes leading up to categorization, they may not be capturing the categorization process itself. For the replication experiment, we observed a similar maximum R^2^ (0.11 versus 0.10 in main dataset), with a peak encoding latency that was slightly earlier (74 ms versus 98 ms, see **Figure 6-1**).

**Figure 6:**
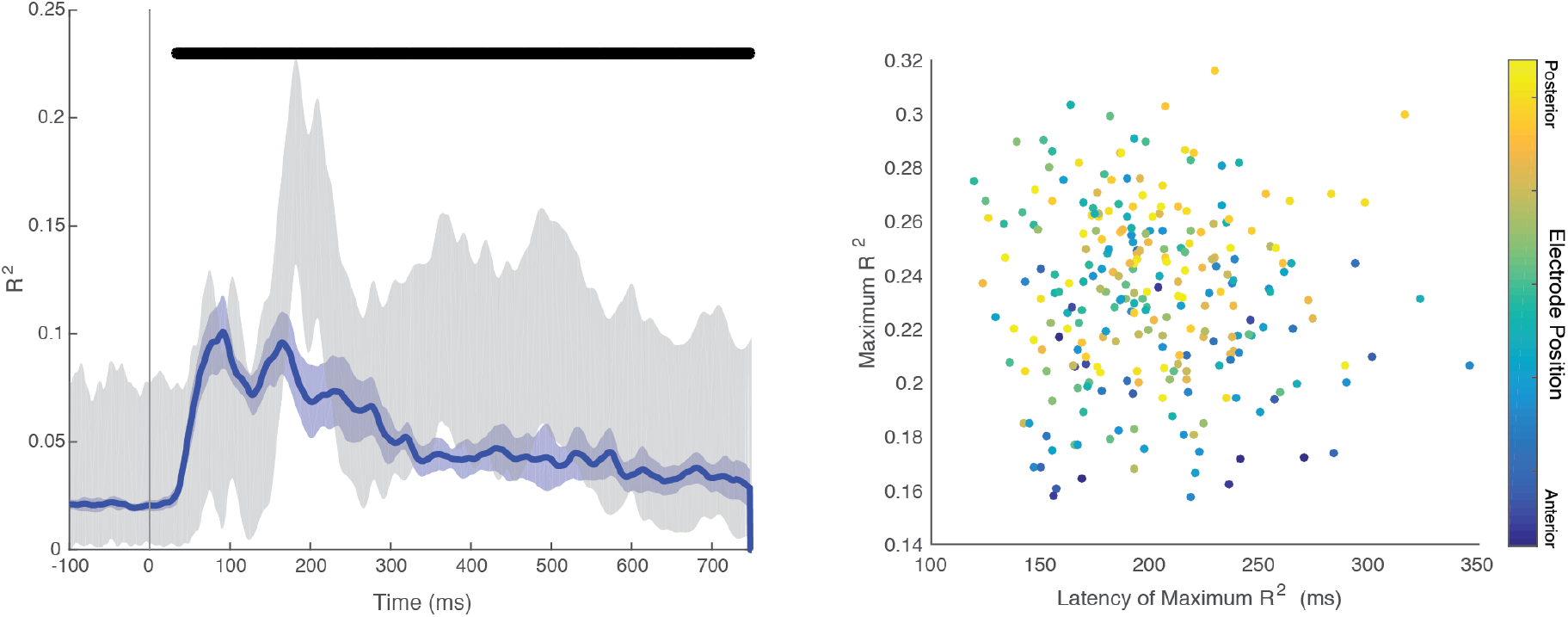
Left: Explained variability (adjusted R^2^) over time. Blue shaded area indicates the 95% confidence interval, thick black bar indicates statistical significance, and gray shaded area represents the noise ceiling of the data. Right: Maximum explained variability (adjusted R^2^) versus latency of maximum R^2^. Each point is an electrode, and color indicates electrode position from anterior to posterior. Data are averaged across participants.

Unlike the case of decoding, we did not observe any relationship between the maximum explained variability in a given electrode and the latency of maximum explained variability (r = −0.04, 95% CI: −0.16-0.08, t(12) < 1), see **Figure 6**. However, as we observed in decoding, there was a reasonably strong spatial relationship between electrode position (anterior to posterior) and maximum R^2^ r = 0.36, 95% CI: 0.25-0.47, t(12) = 3.08, p < 0.01, see **Figure 6**), indicating that these nine models could best explain activity over the posterior electrodes. Similar results were found in the replication experiment (see **Figure 6-2**). Together, these results show that feature encoding is earlier than category decoding, and also primarily driven by posterior electrodes. Our next analyses will examine which specific features predict variability in ERP signals.

#### 3.4.2 Low-level versus high-level features

Among the nine features are those that can be computed directly from image filters (low-level features) as well as those that require human annotation (high-level features). In order to understand the relative contributions of low- and high-level features, we created two new whitened models using a subset of the nine original models. The ‘low-level’ model included all of the filter-based models (Wavelet, Gist, and Texture), and the ‘high-level’ model contained all of the human-in-the-loop models (Objects, Functions, and Attributes). We whitened each model separately and performed the regression analyses on each. Overall, we found that low-level features explained significantly more variability than did the high-level features (0.07 versus 0.03, t(12) = 5.81, p<0.0001, d = 2.24, see **Figure 7**). We observed a similar result in the replication experiment (0.05 versus 0.03, t(14)=30.4, p<1.8e-14, d=12.1, see **Figure 7-1**). When examining the latency of maximum explained variability, low-level models peaked 88 ms after stimulus onset while high-level features peaked 169 ms after stimulus onset (t(12) = 3.18, p<0.005, d = 1.31). Similarly, low-level models peaked 74 ms after stimulus onset in the replication experiment, while high-level models peaked at 159 ms (t(14)=64.7, p<4.8e-19, d=33.4). Thus, low-level features explain more and earlier vERP variability, compared with the high-level features. While this may suggest that high-level features are processed later, it may instead reflect the fact later processes are less time-locked to stimulus onset, and that the increased temporal variability results in a smaller overall effect, or that the neural sources for these features are less accessible at the scalp.

**Figure 7:**
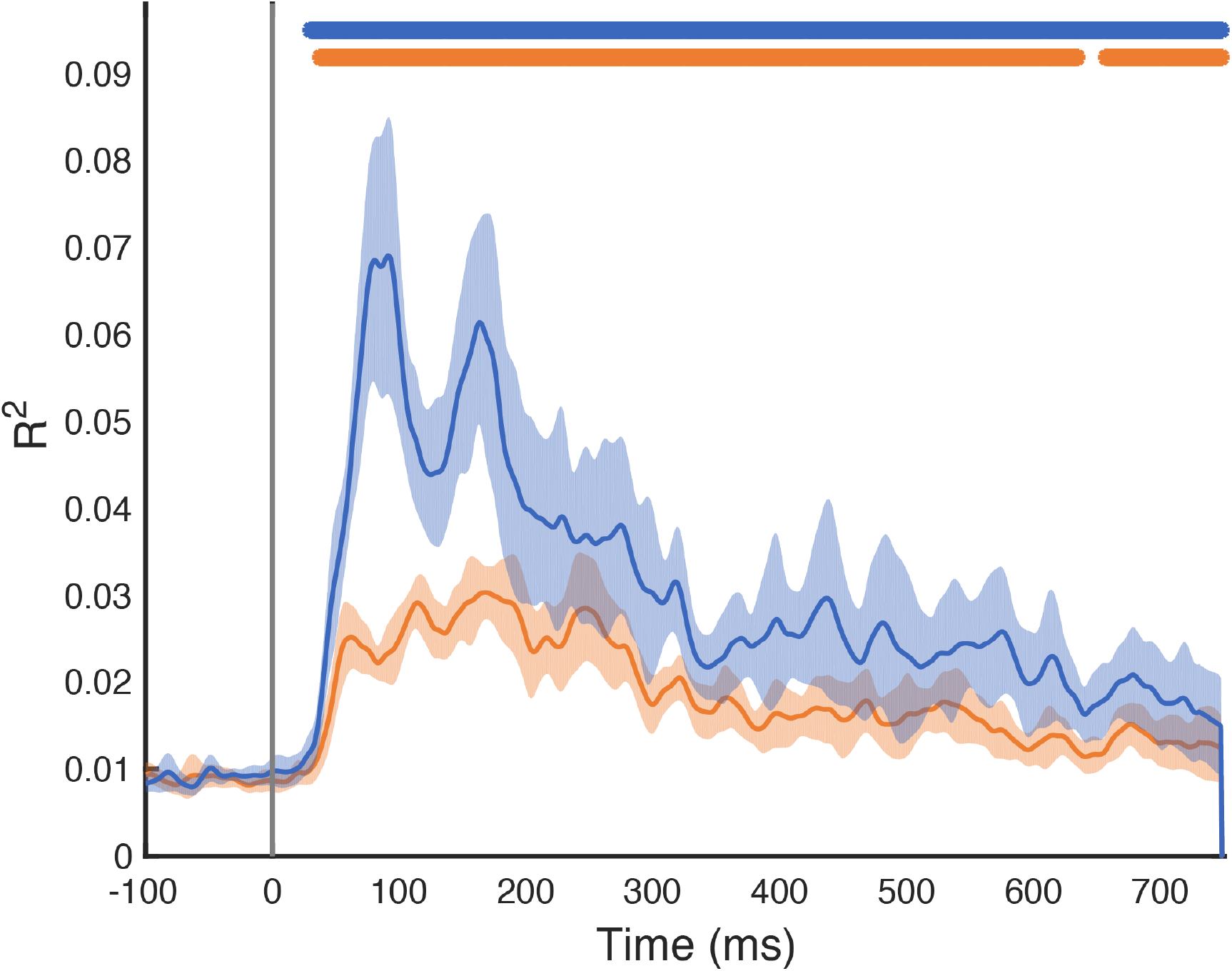
Explained variability in vERP signals for low-level (wavelet, gist, and texture, shown in blue) and high-level (function, object, attribute, shown in orange). Shaded regions represent 95% confidence intervals. Solid lines indicate statistically significant explained variability over baseline (sign permutation test).

#### 3.4.3 Individual Features

The previous analysis showed that low-level features, on average, explained more and earlier variability in vERP responses. Here, we took a closer look by examining the variability explained by each of the nine whitened models individually, see **Figure 8**. **Table 4** shows the maximum r^2^ and the latency of maximum R^2^ for each of the nine feature models. We submitted both of these three measurements to a one-way repeated measures ANOVA with Model (nine features) as the factor. We found that maximum R2 differed significantly across Model (F(8,96) = 7.65, p<7.1e-8, *η*^2^=0.99). Overall, the Gist features explained the most variability in ERPs (R^2^ = 0.026), and explained significantly more variability than all other features except for Wavelet (t(12) = 1.72, p = 0.06), and Conv2 (t(12) < 1). In a similar manner, Conv2 explained more variability than Wavelet (t(12) = 2.22, p < 0.05, d = 0.64), Texture (t(12) = 2.43, p < 0.05, d = 0.98), FC6 (t(12) = 2.34, p < 0.05, d = 0.90), and all of the high-level visual features. Texture had higher explained variance than each of the high-level visual features. We did not observe any significant differences in explained variability among the high-level features. We observed the same main effect of Model on maximum R^2^ in the replication experiment (F(8,112)=5.77, p<3.7e-6, *η*^2^=0.98), and we furthermore observed a high correlation between maximum R2 values between the two experiments (r=0.81, t(7)=3.7, p<0.008), see **Figure 8-1**. Although we observed a numerical tendency for lower-level models to have earlier maximum explained variability, this pattern was not statistically reliable (F(8,96) < 1). This was also observed in the replication experiment (F(8,112)<1).

**Figure 8:**
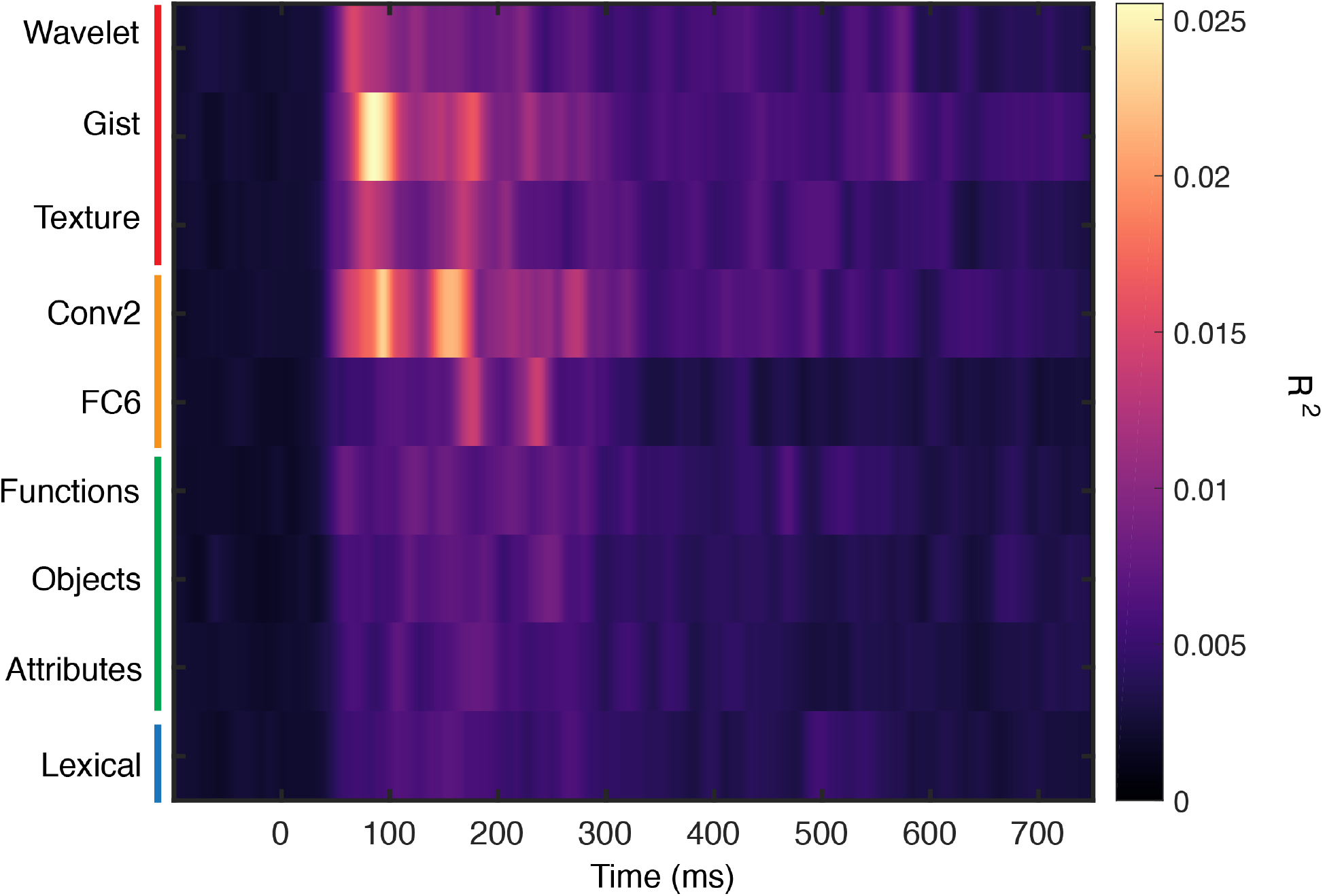
R^2^ values for each of the nine whitened feature models for explaining vERPs, averaged over all electrodes with significant category information.

**Table 4:** Maximum R^2^ and latency of maximum R^2^ for each of the nine encoding models.

| Model | Maximum.R2 | Latency.of.max |
| --- | --- | --- |
| Wavelet | 0.015 | 69 |
| Gist | 0.026 | 85 |
| Texture | 0.014 | 100 |
| Conv2 | 0.023 | 104 |
| FC6 | 0.015 | 205 |
| Function | 0.009 | 135 |
| Object | 0.009 | 238 |
| Attributes | 0.008 | 183 |
| Lexical | 0.007 | 146 |

### 3.5 Linking Encoding and Behavior

#### 3.5.1 Linking vERPs to scene similarity assessment

While the previous encoding analyses examine the extent to which various features explain vERP responses, they do not provide insight into which parts of the responses are behaviorally relevant. To bridge this gap, we first used the behavioral RDM from the unconstrained scene similarity experiment to predict the vERP RDMs. **Figure 9** shows the variability explained over time. We observed an early correspondence of behavioral similarity and neural similarity: the peak R^2^ occurred 219 ms post-stimulus. The mean maximum explained variability was 0.0093 (18.6% of noise ceiling lower bound, versus 0.008 in the replication experiment), with an earlier secondary peak occurring around 175 ms post-stimulus onset. In the replication experiment, only the earlier peak was evident at 178 ms post-image onset (see **Figure 9-1**).

**Figure 9:**
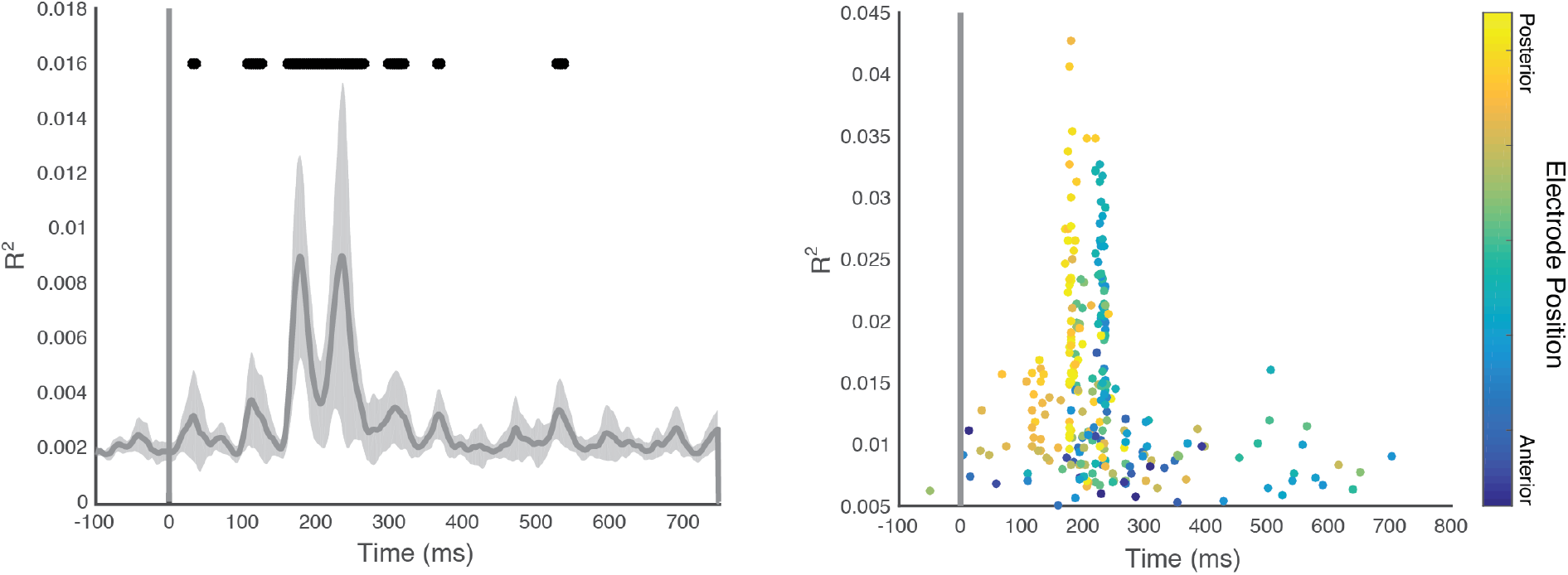
Left: Explained variability (adjusted R^2^) over time for behavioral RDMs for all electrodes with significant category information. Shaded area indicates 95% confidence interval. Right: Maximum explained variability and latency of maximum explained variability for each of the 256 electrodes. Point color indicates electrode position in the net.

In order to determine how each of the task-driven similarity assessments are reflected in the vERPs compared with the unconstrained similarity task examined above, we used each of the five whitened behavioral RDMs from the task-driven similarity experiments as regressors for the vERPs. As shown in **Figure 10**, although using a greater variety of behavioral predictors led to greater explained variability (max of 0.0093 for unconstrained similarity experiment, versus a max of 0.027 for the five task-driven experiments, t(12) = 9.38, p < 3.6e-07, d = 0.91), the time course of the two analyses is strikingly similar, with peaks of explained variability at around 175 ms and 225 ms post-stimulus onset. Similar results were found in the replication experiment (maximum R^2^: 0.027, peak latency: 178 ms, see **Figure 10-1**). As shown in the right-hand panel of **Figure 10**, the earlier peak was driven primarily by posterior electrodes, while the later peak was driven more by anterior electrodes, as was observed in the unconstrained similarity experiment.

**Figure 10:**
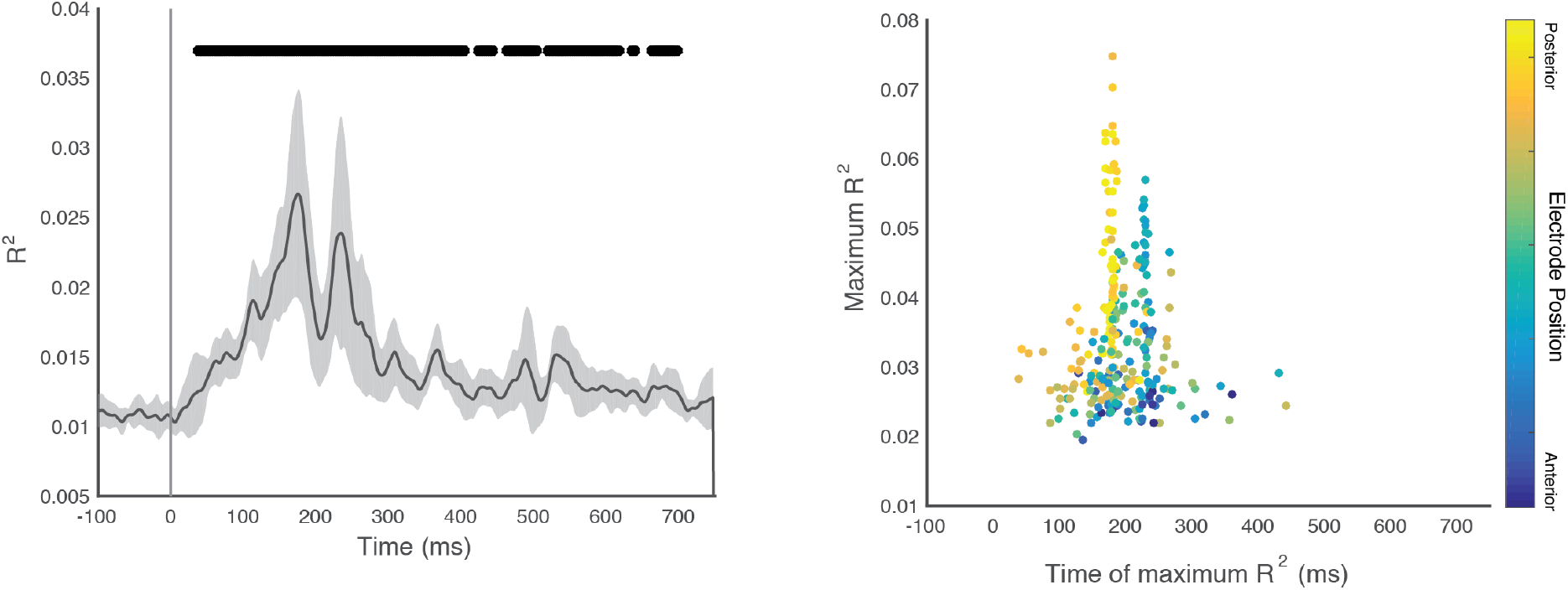
Left: Explained variability (adjusted R^2^) over time for behavioral RDMs for all electrodes with significant category information. Each of the five task-driven experiments was a predictor here. Shaded area indicates 95% confidence interval. Right: Maximum explained variability and latency of maximum explained variability for each of the 256 electrodes. Point color indicates electrode position in the net.

While the previous analysis indicated that all five task-driven similarity experiments explained vERPs in a similar manner as the unconstrained similarity experiment, here we examined each of the five feature-driven similarity experiments individually. As shown in **Figure 11**, each experiment produced a unique profile of explained vERP variability. We performed a one-way ANOVA on the maximum explained variability for each of the five experiments and found that there were significant differences across behavioral experiments in the main EEG experiment (F(4,48) = 3.97, p = 0.0007, *η^2^* = 0.97), as well as the replication experiment (F(4,56)=2.63, p<0.05, *η*^2^ = 0.96, see **Figure 11-1**). Follow-up t-tests revealed that the lexical experiment explained significantly more variability than the orientation (t(12) = 2.47, p = 0.015, d = 0.98), texture (t(12) = 2.02, p = 0.033, d = 0.82), object (t(12) = 2.28, p = 0.021, d = 0.78), and function (t(12) = 2.69, p = 0.0098, d = 0.98) experiments. By contrast, a one-way ANOVA examining differences in peak latency did not reveal any significant differences between the behavioral experiments in either the main EEG experiment (F(4,48) < 1), nor the replication experiment (F(4,56)<1).

**Figure 11:**
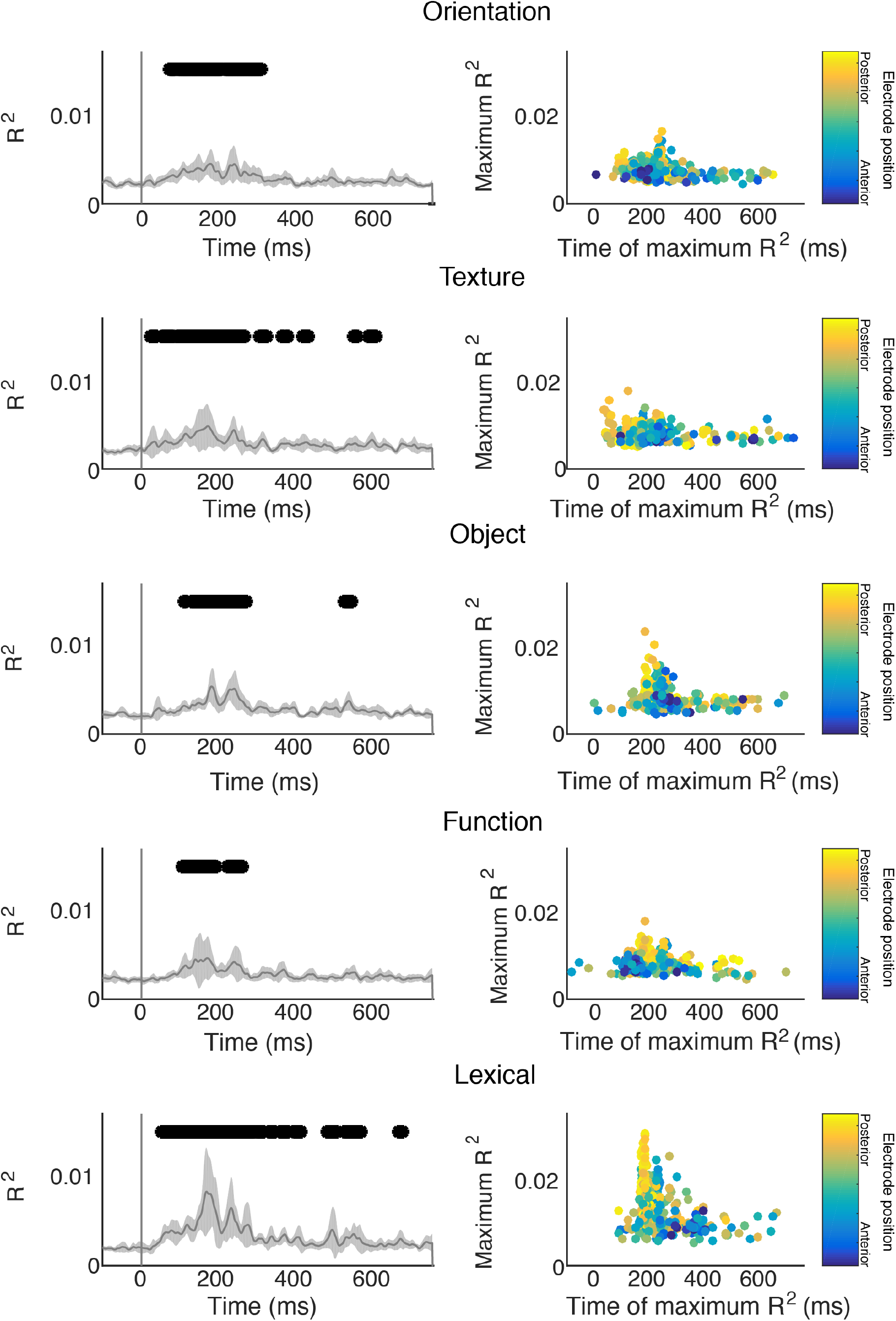
Left panels show explained variability (R^2^) over time of each of the five task-driven similarity experiments for vERPs. Shaded gray regions reflect 95% confidence intervals. Right panels show the relationship between electrode position, maximum R2, and latency of maximum R2 for the same experiments.

#### 3.5.2 Shared R^2^ between scene similarity assessments and features on vERPs

Having established which portions of the vERP response that are linked to scene categorization behavior, we can now examine how vERP variability is shared by each of the nine feature models and the behavioral RDMs for both constrained and unconstrained experiments. This allows us to examine when and how individual features explain category-relevant portions of the vERP data. Beginning with the unconstrained similarity experiment, **Figure 12** shows time-resolved plots of shared R^2^ for all feature models and unconstrained similarity judgment over the time epoch. **Table 5** shows maximum shared R^2^ and latency of maximum shared R^2^ values for each of the nine whitened features. We observed a significant difference between features in the maximum shared variability with unconstrained similarity judgments (F(8,96) = 17.91, p=4.3e-16, *η*^2^ = 0.99 in the main experiment and F(8,112) = 15.44, p = 4.11e-15, *η*^2^ = 0.99 in the replication experiment). The maximum shared variability ranged from 0.00008 for the Conv2 model to 0.0049 for the FC6 model and was higher for high-level models than low-level models (0.0018 versus 0.002, t(12) = 4.82, p = 0.002, d = 1.72 in the main experiment; 0.002 versus 0.0002, t(14)=5.93, p=1.84e-05, d=1.65 in the replication experiment). The latency of maximum shared variance ranged from 168 ms post-stimulus for the Conv2 model to 235 ms post-stimulus for Objects. These differences were not statistically reliable when considering all nine models (F(8,96)<1 in the main experiment and F(8,112)<1 in the replication experiment), nor when comparing the three low-level features (Wavelet, Gist, and Texture) to the three high-level models (Functions, Objects, and Attributes), (176 versus 200 ms, t(12)<1 in the main experiment; 155 versus 201 ms, t(14)<1 in the replication experiment). However, when comparing the latency of maximum shared variability to the latency of maximum non-shared variability, vERP variability that was shared with behavior was later on average than non-shared variability (193 ms post-image onset versus 144 ms, t(12) = 12.7, p = 2.4e-08, d = 4.98 for the main dataset, 158 ms versus 138 ms, t(14) = 7.4, p = 3.28e-6, d = 2.07 for the replication experiment). Therefore, although low-level models explain more and earlier variability in vERPs overall, they explain less behaviorally-relevant vERP information when compared with the high-level features, and the time course of behaviorally-relevant shared variability is slower overall than feature processing that is not associated with behavior.

**Figure 12:**
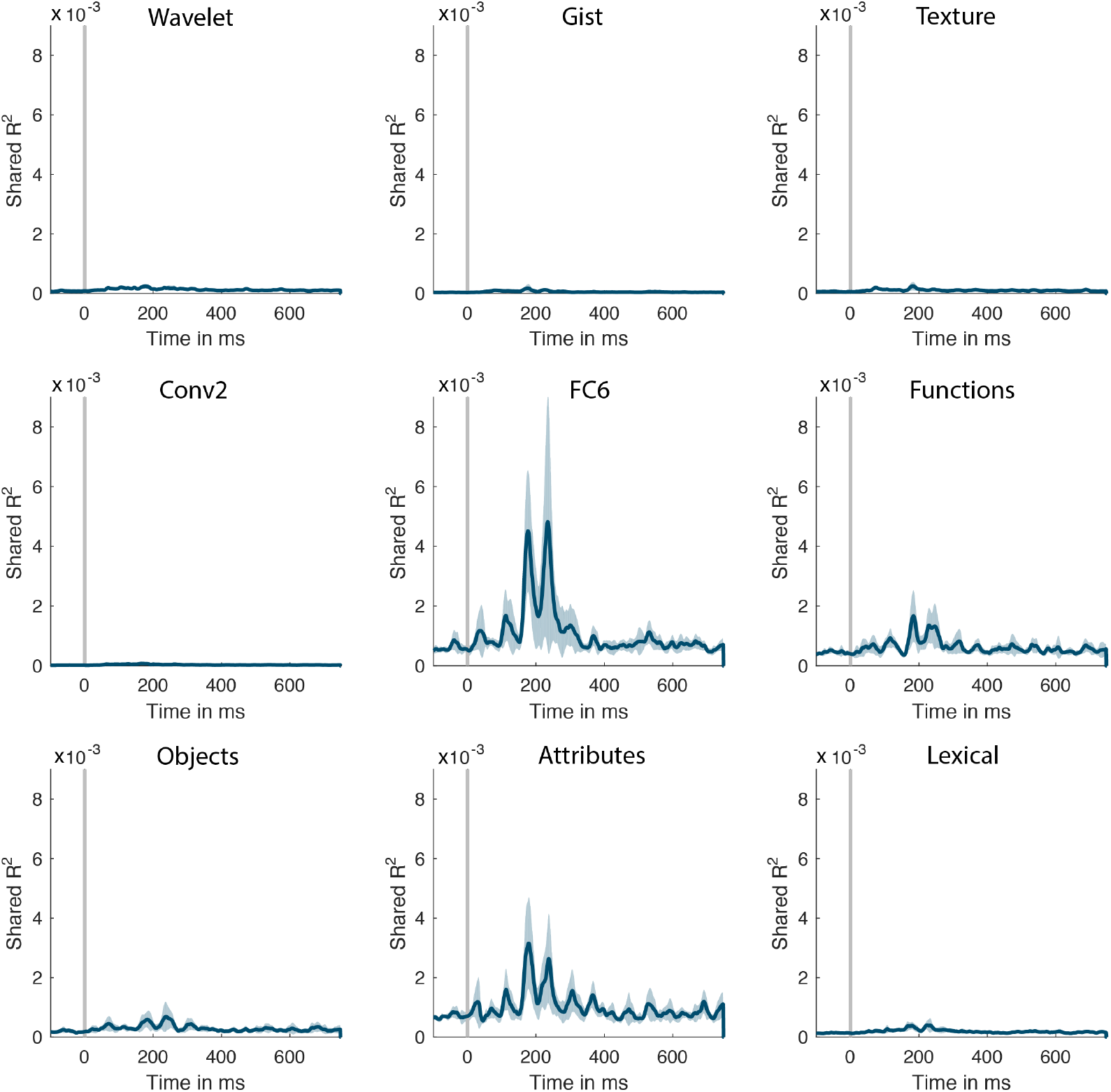
Shared explained variance between each of the nine feature models and the RDM for unconstrained scene similarity judgments over time.

**Table 5:**
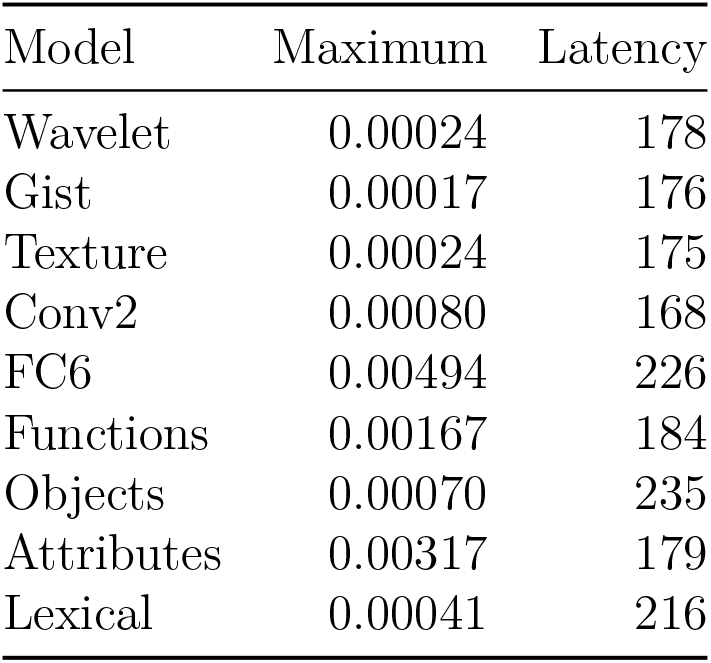
Statistics of shared variability between features, unconstrained behavior, and the nine feature models. Displayed are maximum R^2^ and latency of maximum R^2^ for each of the nine encoding models.

Finally, we followed the same procedure for each of the five task-specific experiments. The aggregate results are shown in **Figure 13**. We found that there was no significant difference in maximum shared variability across the five experiments (F(4,48) = 2.18, p = 0.09, but found a marginal effect in the replication experiment (F(4,56) = 2.81, p = 0.03, *η*^2^ = 0.84), see **Figure 13-1**). However, we observed a significant effect of Feature (F(8,96) = 21.67, p = 1.78e-18, *η*^2^ = 0.99 in the main experiment, and F(8,112) = 13.3, p = 2.1e-13, *η*^2^ = 0.99 in the replication experiment). Following up on this result, we found that high-level features shared significantly more behaviorally-relevant variance with vERPs than low-level features (0.0007 versus 0.0003, t(12) = 5.08, p = 0.00014, d = 1.67 in the main experiment and 0.0008 versus 0.0003, t(14) = 7.49, p = 1.46e-6, d = 2.04 in the replication study). Finally, we observed a significant interaction between Feature and Experiment: F(32, 384) = 7.92, p = 1.52e-26, *η*^2^ = 0.98 in the main experiment and F(32, 488) = 11.11, p = 1.8e-39, *η*^2^ = 0.99 in the replication experiment). Interestingly, this was driven by the fact that the orientation experiment had higher shared variability with low-level features compared with high-level (t(12) = 3.21, p = 0.003, d = 1.37 in the main experiment and t(14) = 2.67, p = 0.009, d = 0.83 in the replication study), while the opposite pattern was found for the four other experiments. Therefore, although high-level features seem to be more behaviorally-relevant overall, this can change when the task demands that observers attend to low-level scene properties. We observed no significant effects of Experiment (F(4,48)<1 and F(4,56)<1) or Feature (F(8,96)<1 and F(8,112)<1) on the latency of maximum shared variance, nor a significant interaction between these two factors (F(32,384)<1 and F(32,448)<1).

**Figure 13:**
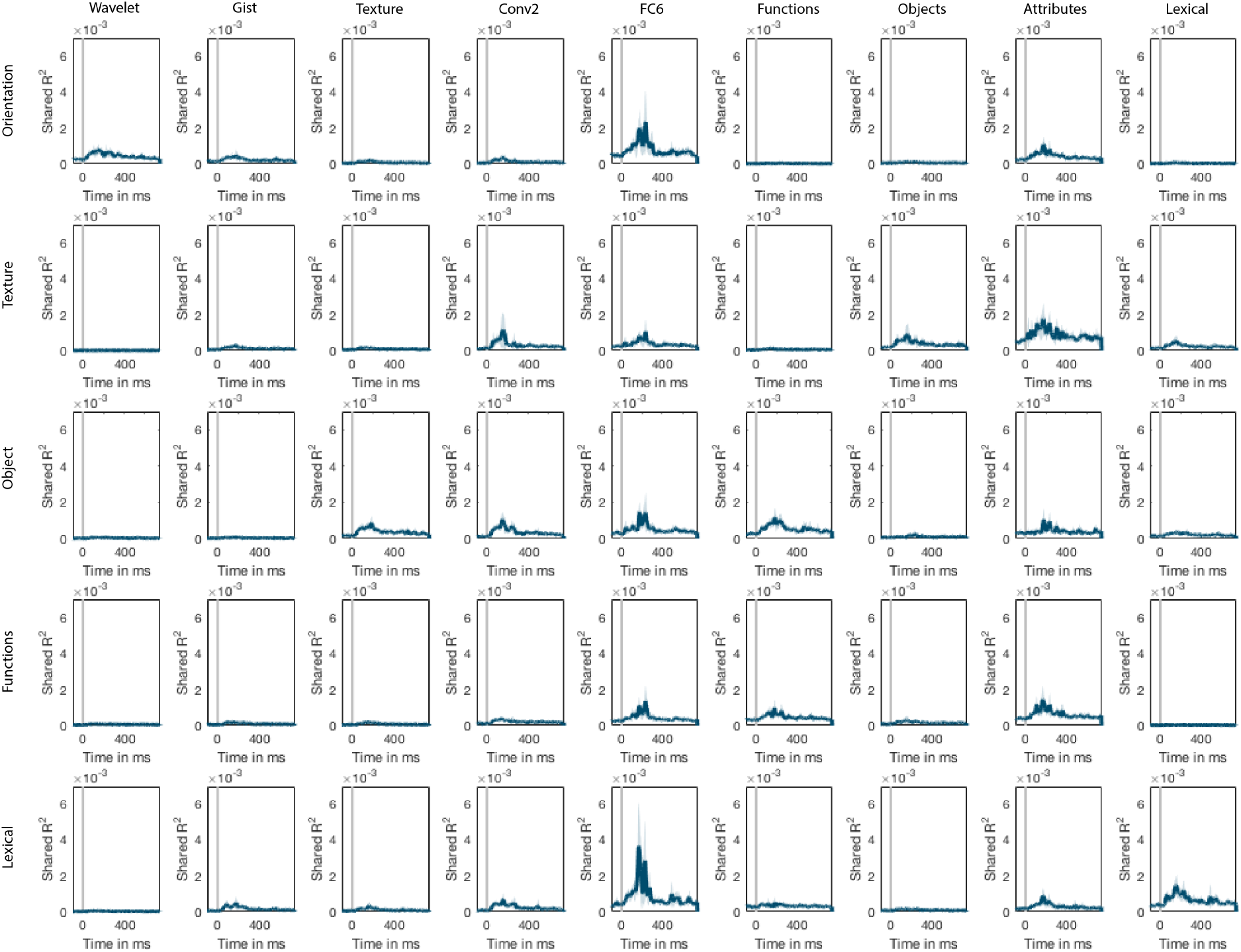
Shared variability of each feature (columns) with vERPs across each of the five task-driven experiments (rows).

Thus, compared with the unconstrained similarity experiment, we can see that changing the behavioral task changes the amount of shared vERP variability with features that are associated with the task. Specifically, although the wavelet features share little behaviorally-relevant vERP variability in most experiments, this was not the case for the orientation task where these low-level features were task-relevant. Similarly, the lexical features shared more behaviorally-relevant vERP variability in the lexical experiment compared with the others.

## 4. Discussion

Visual categorization is rapid and seemingly effortless. However, the initial visual input must be transformed into alternative representations that allow categories to be easily distinguished from one another (DiCarlo & Cox, 2007). Here, we sought to understand the visual processing stages required to transform the retinal image into a semantically rich categorical representation. We first verified the utility of a variety of popular visual and conceptual feature models for scene categorization (Section 3.1). Using time-resolved decoding, we assessed the amount of category-related information available in vERPs (Section 3.3). We then applied a whitening transformation to the feature models and assessed their utility for explaining vERPs (Section 3.4). Critically, we then assessed the shared variability between features and behavior for explaining vERPs (Section 3.5). By combining encoding, decoding, and behavioral assessments, we can link neural activity to feature spaces (encoding), as well to the time course of category information (decoding), and to the internal representations that guide scene categorization behavior.

Our decoding results revealed that decodable scene category information peaked between 150 and 200 ms after image onset and persisted across the trial epoch. These values are consistent with previous M/EEG studies of object- and scene categorization (Bankson, Hebart, Groen, & Baker, 2018; Carlson, Tovar, Alink, & Kriegeskorte, 2013; Cichy, Pantazis, & Oliva, 2014; Clarke, Taylor, Devereux, Randall, & Tyler, 2013; Ramkumar et al., 2016). While earlier decoding has been reported for image exemplars (~100 ms, (Carlson et al., 2013; Cichy et al., 2014)), it has remained unclear whether this performance reflects image identity per se, or the lower-level visual features that are associated with that exemplar.

In contrast to previous work, we have tested an extensive set of features ranging from low-level filter outputs to high-level conceptual features that require extensive human annotation. Each of the nine features used here has been implicated in scene categorization. Nearly all have been shown to be computationally sufficient for categorization, and many have striking correlations with brain activity and behavior. However, because these models are often studied in isolation, and because they are correlated with one another, it has been difficult to assess the *independent* contributions of each. Here, we employed a whitening transformation to the input feature RDMs in order to decorrelate the feature spaces. Although there is increasing understanding of the need to partition explained variability for correlated inputs (Bankson et al., 2018; Greene et al., 2016; Groen et al., 2018; Lescroart, Stansbury, & Gallant, 2015), it is difficult to do this for a large number of input models. We side-stepped these issues by whitening the features before fitting models. There are many whitening transforms, and we chose the ZCA algorithm because it has been shown to provide outputs that are best correlated with the original inputs (Kessy et al., 2017). While this generally held true for the nine models used here, it should be noted that the gist features (Oliva & Torralba, 2001) were an exception (see **Figure 1**). Therefore, we have refrained from strongly interpreting results for that model, particularly the observation that this feature was not significantly predictive of the behavioral RDM (see **Table 1**) given previous reports that gist features can strongly influence categorization behavior (M. Greene & Oliva, 2009), vERPs (Hansen, Noesen, Nador, & Harel, 2018), MEG patterns (Ramkumar et al., 2016), and fMRI activation patterns (Watson et al., 2017, 2014).

The current results demonstrate that low-level visual features explained the earliest variability in vERPs (~90 ms post-image onset). High-level visual features had the highest explained variability 80 ms later (~170 ms), similar to the time course of predicting vERPs with the unconstrained behavioral data (~175 ms), or the aggregate of all five scene similarity tasks (**Figure 10** and **Figure 11**). Further, the average peak decoding accuracy was observed ~200 ms, and peak shared variability for each feature with behavior also ranged between ~170-230 ms post-image onset. Together, this suggests a progression to categorization that proceeds from low-level to high-level features.

The observed time course of semantic processing may seem faster than previously characterized ERPs such as the N400 (Kutas & Federmeier, 2000). Indeed, violations of scene-object context have been observed in the 250-350 ms post-image window (Mudrik, Shalgi, Lamy, & Deouell, 2014), as well as in the classical N400 window (Ganis & Kutas, 2003; Võ & Wolfe, 2013). However, recent decoding results have shown that these two windows contain similar image information (Draschkow, Heikel, Võ, Fiebach, & Sassenhagen, 2018) and may be reflecting similar neural processes. It is worth noting that the time course of these ERPs reflects an upper bound to the time course of semantic processing, and that the current encoding and decoding techniques may reveal the processes themselves while the ERPs reflect the outcomes. However, many leading theories of the N400 characterize it as a full contextual evaluation of the stimulus (Kutas & Federmeier, 2000), and this may require a full categorical representation before this evaluation can take place. Consistent with this idea is our observation that behaviorally-relevant vERP variance was shared after the decoding peak, particularly for high-level features (see **Figures 6, 7, 7**, and **12**), suggesting that they contributed both to category representations and post-categorization processing.

When considering all nine models together, the explained variability for vERPs was largely within the noise ceiling of the data, indicating that these models’ predictive power has been maximized, given the noise in the data. We observed two distinct R^2^ peaks, one around 100 ms after image onset, and the other around 75 ms later (see **Figure 6**). While low-level features contributed to both peaks (see **Figure 7**), most of the contribution from high-level models was during this later period (see **Table 4**). These results are consistent with other reports of low-level feature encoding (Groen et al., 2012; Hansen et al., 2011, 2012). Critically, it is only this later peak that is correlated with scene categorization behavior (see **Figure 12**). Therefore, although low-level features are critical for subsequent categorization, they do not themselves enable categorization, counter to views of scene categorization being largely associated with low-level features (Kaping, Tzvetanov, & Treue, 2007; Scholte, Ghebreab, Waldorp, Smeulders, & Lamme, 2009; Torralba & Oliva, 2003).

Our results are congruent with previous ERP studies that have shown that evoked responses earlier than ~150 ms post-stimulus onset are not correlated with behavioral measurements (Johnson & Olshausen, 2003; Philiastides & Sajda, 2006; VanRullen & Thorpe, 2001). However, our results extend those previous findings by allowing us to make inferences about the visual and conceptual features that are associated with those behaviorally-relevant neural signals. Specifically, our results indicate that high-level features share more with scene similarity responses than do low-level features. Specifically, the higher-level features from the dCNN (FC6) and the attribute model shared the most variability with vERPs and unconstrained similarity assessment. Deep CNN models are optimized for categorization, and the representations in their upper layers reflect this fact. The attribute model, as discussed in Section 3.1 is a heterogeneous model reflecting human annotations of affordances, surfaces, materials and spatial properties. Thus, the whitened attribute model likely reflects aspects of texture, objects, and affordances that are not captured in those individual models.

While much of the scene understanding literature focuses on scene categorization as an “end point” of the visual recognition process, it is important to recognize that perception is an ongoing process without a strict end (Groen et al., 2017; Malcolm et al., 2016). However, categories are highly linked to other behavioral tasks, including object detection (Davenport & Potter, 2004), visual search (Torralba, Oliva, Castelhano, & Henderson, 2006), and navigation (Bonner & Epstein, 2018). Therefore, we have utilized two types of behavioral tasks: an unconstrained scene similarity assessment task that has previously been shown to reveal hierarchical category representations (Zheng et al., 2019), and a set of five tasks that ask observers to assess scene similarity with respect to one of five features that were designed to have observers attend to low- (orientation), mid- (texture), and high-level features (objects, functions, and lexical). We have shown that changing the task changes the shared variability between vERPs and features (see **Figure 13**). Specifically, although some features share little variability with the scene categorization, they seem to be used when the task demands it. This is striking because the participants in the behavioral experiments were independent of those in the EEG experiment. We are currently extending this paradigm to change the observers’ task during EEG recording (Hansen & Greene, 2019).

By using a combination of encoding and decoding approaches on high-density EEG data, we have shown that the visual processes leading up to scene categorization follow a progression from low- to high-level feature processing from occipital through ventral and medial temporal cortices in the first 200 ms after scene onset. While low-level features explain more vERP variability overall, they tend not to share variability with behavioral tasks, except for when those features are task-relevant. Altogether, these results call into question models of scene categorization that are based solely on low-level features, and further highlight the flexible nature of the categorization process.

## Acknowledgments

National Science Foundation grant (1736394) to MRG and BCH. James S. McDonnell Foundation grant (220020430) to BCH.

## References

Bankson, B. B., Hebart, M. N., Groen, I. I. A., & Baker, C. I. (2018). The temporal evolution of conceptual object representations revealed through models of behavior, semantics and deep neural networks. NeuroImage, 178, 172–182.

Bastin, J., Vidal, J. R., Bouvier, S., Perrone-Bertolotti, M., Bénis, D., Kahane, P., David, O., et al. (2013). Temporal Components in the Parahippocampal Place Area Revealed by Human Intracerebral Recordings. The Journal of Neuroscience, 33 (24), 10123–10131.

Bell, A. J., & Sejnowski, T. J. (1997). The “independent components” of natural scenes are edge filters. Vision Research, 37 (23), 3327–3338.

Biederman, I. (1981). On the semantics of a glance at a scene. In Perceptual Organization. New Jersey: Hillsdale.

Bonner, M. F., & Epstein, R. A. (2018). Computational mechanisms underlying cortical responses to the affordance properties of visual scenes. PLOS Computational Biology, 14 (4), e1006111.

Bruss, F. T. (2000). Sum the Odds to One and Stop. The Annals of Probability, 28 (3), 1384–1391.

Cadieu, C. F., Hong, H., Yamins, D. L. K., Pinto, N., Ardila, D., Solomon, E. A., Majaj, N. J., et al. (2014). Deep Neural Networks Rival the Representation of Primate IT Cortex for Core Visual Object Recognition. PLOS Computational Biology, 10 (12), e1003963.

Cant, J. S., & Goodale, M. A. (2011). Scratching Beneath the Surface: New Insights into the Functional Properties of the Lateral Occipital Area and Parahippocampal Place Area. The Journal of Neuroscience, 31 (22), 8248–8258.

Carandini, M., Demb, J. B., Mante, V., Tolhurst, D. J., Dan, Y., Olshausen, B. A., Gallant, J. L., et al. (2005). Do We Know What the Early Visual System Does? Journal of Neuroscience, 25 (46), 10577–10597.

Carlson, T., Tovar, D. A., Alink, A., & Kriegeskorte, N. (2013). Representational dynamics of object vision: The first 1000 ms. Journal of Vision, 13 (10), 1–1.

Cichy, R. M., Khosla, A., Pantazis, D., & Oliva, A. (2017). Dynamics of scene representations in the human brain revealed by magnetoencephalography and deep neural networks. NeuroImage, 153, 346–358.

Cichy, R. M., Khosla, A., Pantazis, D., Torralba, A., & Oliva, A. (2016). Comparison of deep neural networks to spatio-temporal cortical dynamics of human visual object recognition reveals hierarchical correspondence. Scientific Reports, 6, 27755.

Cichy, R. M., Pantazis, D., & Oliva, A. (2014). Resolving human object recognition in space and time. Nature Neuroscience, 17 (3), 455–462.

Clarke, A., Taylor, K. I., Devereux, B., Randall, B., & Tyler, L. K. (2013). From perception to conception: How meaningful objects are processed over time. Cerebral cortex (New York, N.Y.: 1991), 23 (1), 187–197.

Davenport, J., & Potter, M. C. (2004). Scene consistency in object and background perception. Psychological Science, 15 (8), 559–564.

Delorme, A., & Makeig, S. (2004). EEGLAB: An open source toolbox for analysis of singletrial EEG dynamics including independent component analysis. Journal of Neuroscience Methods, 134 (1), 9–21.

DiCarlo, J. J., & Cox, D. D. (2007). Untangling invariant object recognition. Trends in Cognitive Sciences, 11 (8), 333–341.

Draschkow, D., Heikel, E., Võ, M. L. H., Fiebach, C. J., & Sassenhagen, J. (2018). No evidence from MVPA for different processes underlying the N300 and N400 incongruity effects in object-scene processing. Neuropsychologia, 120, 9–17.

Edelman, S. (1998). Representation is representation of similarities. The Behavioral and Brain Sciences, 21 (4), 449–467; discussion 467–498.

Fei-Fei, L., & Perona, P. (2005). A Bayesian Hierarchical Model for Learning Natural Scene Categories. In Proceedings of the 2005 IEEE Computer Society Conference on Computer Vision and Pattern Recognition (CVPR’05) - Volume 2 - Volume 02 (pp. 524–531). IEEE Computer Society.

Field, D. J. (1987). Relations between the statistics of natural images and the response properties of cortical cells. Journal of the Optical Society of America, 4 (12), 2379–2394.

Freeman, J., & Simoncelli, E. P. (2011). Metamers of the ventral stream. Nature Neuroscience, 14 (9), 1195–1201.

Ganis, G., & Kutas, M. (2003). An electrophysiological study of scene effects on object identification. Cognitive Brain Research, 16 (2), 123–144.

Gärdenfors, P. (2004). Conceptual Spaces: The Geometry of Thought. MIT Press.

Greene, M., & Oliva, A. (2009). Recognition of natural scenes from global properties: Seeing the forest without representing the trees. Cognitive Psychology, 58 (2), 137–176.

Greene, M. R. (2013). Statistics of high-level scene context. Frontiers in Perception Science, 4, 777.

Greene, M. R., Baldassano, C., Esteva, A., Beck, D. M., & Fei-Fei, L. (2016). Visual scenes are categorized by function. Journal of Experimental Psychology. General, 145 (1), 82–94.

Greene, M. R., & Hansen, B. C. (2018). Shared spatiotemporal category representations in biological and artificial deep neural networks. PLOS Computational Biology, 14 (7), e1006327.

Greene, M. R., & Oliva, A. (2009). The Briefest of Glances: The Time Course of Natural Scene Understanding. Psychological Science, 20, 464–472.

Groen, I. I. A., Ghebreab, S., Lamme, V. A. F., & Scholte, H. S. (2012). Spatially Pooled Contrast Responses Predict Neural and Perceptual Similarity of Naturalistic Image Categories. PLoS Comput Biol, 8 (10), e1002726.

Groen, I. I. A., Silson, E. H., & Baker, C. I. (2017). Contributions of low-and high-level properties to neural processing of visual scenes in the human brain. Phil. Trans. R. Soc. B, 372 (1714), 20160102.

Groen, I. I., Greene, M. R., Baldassano, C., Fei-Fei, L., Beck, D. M., & Baker, C. I. (2018). Distinct contributions of functional and deep neural network features to representational similarity of scenes in human brain and behavior. eLife, 7.

Güçlü, U., & Gerven, M. A. J. van. (2015). Deep Neural Networks Reveal a Gradient in the Complexity of Neural Representations across the Ventral Stream. The Journal of Neuroscience, 35 (27), 10005–10014.

Hansen, B. C., & Essock, E. A. (2004). A horizontal bias in human visual processing of orientation and its correspondence to the structural components of natural scenes. Journal of Vision, 4 (12), 5–5.

Hansen, B. C., & Greene, M. R. (2019). Task demands flexibly change the dynamics of feature use during scene processing. Journal of Vision.

Hansen, B. C., Jacques, T., Johnson, A. P., & Ellemberg, D. (2011). From spatial frequency contrast to edge preponderance: The differential modulation of early visual evoked potentials by natural scene stimuli. Visual Neuroscience, 28 (3), 221–237.

Hansen, B. C., Johnson, A. P., & Ellemberg, D. (2012). Different spatial frequency bands selectively signal for natural image statistics in the early visual system. Journal of Neurophysiology, 108 (8), 2160–2172.

Hansen, B. C., & Loschky, L. C. (2013). The contribution of amplitude and phase spectra-defined scene statistics to the masking of rapid scene categorization. Journal of Vision, 13 (13), 21–21.

Hansen, N. E., Noesen, B. T., Nador, J. D., & Harel, A. (2018). The influence of behavioral relevance on the processing of global scene properties: An ERP study. Neuropsychologia, 114, 168–180.

Harel, A., Kravitz, D. J., & Baker, C. I. (2012). Deconstructing Visual Scenes in Cortex: Gradients of Object and Spatial Layout Information. Cerebral Cortex.

Isik, L., Meyers, E. M., Leibo, J. Z., & Poggio, T. (2014). The dynamics of invariant object recognition in the human visual system. Journal of Neurophysiology, 111 (1), 91–102.

Jia, Y., Shelhamer, E., Donahue, J., Karayev, S., Long, J., Girshick, R., Guadarrama, S., et al. (2014). Caffe: Convolutional Architecture for Fast Feature Embedding. In Proceedings of the 22Nd ACM International Conference on Multimedia, MM ’14 (pp. 675–678). New York, NY, USA: ACM.

Johnson, J. S., & Olshausen, B. A. (2003). Timecourse of neural signatures of object recognition. Journal of Vision, 3 (7).

Kaping, D. [., Tzvetanov, T. [., & Treue, S. [. (2007). Adaptation to statistical properties of visual scenes biases rapid categorization. Visual Cognition, 15, 12–19.

Kessy, A., Lewin, A., & Strimmer, K. (2017). Optimal Whitening and Decorrelation. The American Statistician, 0 (0), 1–6.

Khaligh-Razavi, S.-M., & Kriegeskorte, N. (2014). Deep Supervised, but Not Unsupervised, Models May Explain IT Cortical Representation. PLOS Comput Biol, 10 (11), e1003915.

Kriegeskorte, N., Mur, M., Ruff, D. A., Kiani, R., Bodurka, J., Esteky, H., Tanaka, K., et al. (2008). Matching Categorical Object Representations in Inferior Temporal Cortex of Man and Monkey. Neuron, 60 (6), 1126–1141.

Krizhevsky, A., Sutskever, I., & Hinton, G. E. (2012). ImageNet Classification with Deep Convolutional Neural Networks. In F. Pereira, C. J. C. Burges, L. Bottou, & K. Q. Weinberger (Eds.), Advances in Neural Information Processing Systems 25 (pp. 1097–1105). Curran Associates, Inc.

Kubilius, J., Bracci, S., & Beeck, H. P. O. de. (2016). Deep Neural Networks as a Computational Model for Human Shape Sensitivity. PLOS Comput Biol, 12 (4), e1004896.

Kuo, C.-C., Luu, P., Morgan, K. K., Dow, M., Davey, C., Song, J., Malony, A. D., et al. (2014). Localizing Movement-Related Primary Sensorimotor Cortices with Multi-Band EEG Frequency Changes and Functional MRI. PLOS ONE, 9 (11), e112103.

Kutas, M., & Federmeier, K. D. (2000). Electrophysiology reveals semantic memory use in language comprehension. Trends in Cognitive Sciences, 4 (12), 463–470.

Lazebnik, S., Schmid, C., & Ponce, J. (2006). Beyond Bags of Features: Spatial Pyramid Matching for Recognizing Natural Scene Categories. In (Vol. 2, pp. 2169–2178). IEEE.

Lescroart, M. D., Stansbury, D. E., & Gallant, J. L. (2015). Fourier power, subjective distance, and object categories all provide plausible models of BOLD responses in scene-selective visual areas. Frontiers in Computational Neuroscience, 9, 135.

Li, K., Papademetris, X., & Tucker, D. M. (2016). BrainK for Structural Image Processing: Creating Electrical Models of the Human Head. Computational Intelligence and Neuroscience, 2016, e1349851.

Lowe, M. X., Rajsic, J., Gallivan, J. P., Ferber, S., & Cant, J. S. (2017). Neural representation of geometry and surface properties in object and scene perception. NeuroImage, 157, 586–597.

Luck, S. J. (2005). An introduction to the event-related potential technique. MIT Press.

MacEvoy, S. P., & Epstein, R. A. (2011). Constructing scenes from objects in human occipitotemporal cortex. Nature Neuroscience, 14 (10), 1323–1329.

Malcolm, G. L., Groen, I. I. A., & Baker, C. I. (2016). Making Sense of Real-World Scenes. Trends in Cognitive Sciences, 20 (11), 843–856.

Miller, G. A. (1995). WordNet: A lexical database for English. Commun. ACM, 38 (11), 39–41.

Mudrik, L., Shalgi, S., Lamy, D., & Deouell, L. Y. (2014). Synchronous contextual irregularities affect early scene processing: Replication and extension. Neuropsychologia, 56, 447–458.

Neely, J. H. (1977). Semantic priming and retrieval from lexical memory: Roles of inhibitionless spreading activation and limited-capacity attention. Journal of Experimental Psychology: General, 106 (3), 226–254.

Nili, H., Wingfield, C., Walther, A., Su, L., Marslen-Wilson, W., & Kriegeskorte, N. (2014). A Toolbox for Representational Similarity Analysis. PLOS Computational Biology, 10 (4), e1003553.

Oliva, A., & Schyns, P. G. (2000). Diagnostic Colors Mediate Scene Recognition. Cognitive Psychology, 41 (2), 176–210.

Oliva, A., & Torralba, A. (2001). Modeling the Shape of the Scene: A Holistic Representation of the Spatial Envelope. International Journal of Computer Vision, 42 (3), 145–175.

Patterson, G., Xu, C., Su, H., & Hays, J. (2014). The SUN Attribute Database: Beyond Categories for Deeper Scene Understanding. International Journal of Computer Vision, 108 (1-2), 59–81.

Pedersen, T., Patwardhan, S., & Michelizzi, J. (2004). WordNet::Similarity: Measuring the Relatedness of Concepts. In Demonstration Papers at HLT-NAACL 2004, HLT-NAACLDemonstrations ’04 (pp. 38–41). Stroudsburg, PA, USA: Association for Computational Linguistics.

Philiastides, M. G., & Sajda, P. (2006). Temporal Characterization of the Neural Correlates of Perceptual Decision Making in the Human Brain. Cerebral Cortex, 16 (4), 509–518.

Portilla, J., & Simoncelli, E. (2000). A parametric texture model based on joint statistics of complex wavelet coefficients. International Journal of Computer Vision, 40 (1), 49–71.

Potter, M. C., Wyble, B., Hagmann, C. E., & McCourt, E. S. (2014). Detecting meaning in RSVP at 13 ms per picture. Attention, Perception, & Psychophysics, 1–10.

Ramkumar, P., Hansen, B. C., Pannasch, S., & Loschky, L. C. (2016). Visual information representation and rapid-scene categorization are simultaneous across cortex: An MEG study. NeuroImage, 134, 295–304.

Renninger, L. W., & Malik, J. (2004). When is scene indentification just texture recognition? Vision Research, 44, 2301–2311.

Russell, B., Torralba, A., Murphy, K., & Freeman, W. (2008). LabelMe: A Database and Web-Based Tool for Image Annotation. International Journal of Computer Vision, 77 (1), 157–173.

Scholte, H. S., Ghebreab, S., Waldorp, L., Smeulders, A. W. M., & Lamme, V. A. F. (2009). Brain responses strongly correlate with Weibull image statistics when processing natural images. Journal of Vision, 9 (4).

Sermanet, P., Eigen, D., Zhang, X., Mathieu, M., Fergus, R., & LeCun, Y. (2013). OverFeat: Integrated Recognition, Localization and Detection using Convolutional Networks. arXiv:1312.6229 [cs]. Retrieved from http://arxiv.org/abs/1312.6229

Simoncelli, E. P., & Freeman, W. T. (1995). The steerable pyramid: A flexible architecture for multi-scale derivative computation. In Proceedings., International Conference on Image Processing (Vol. 3, pp. 444–447 vol.3).

Song, J., Davey, C., Poulsen, C., Luu, P., Turovets, S., Anderson, E., Li, K., et al. (2015). EEG source localization: Sensor density and head surface coverage. Journal of Neuroscience Methods, 256, 9–21.

Thorpe, S., Fize, D., & Marlot, C. (1996). Speed of processing in the human visual system. Nature, 381, 520–522.

Torralba, A., & Oliva, A. (2003). Statistics of natural image categories. Network (Bristol, England), 14 (3), 391–412.

Torralba, A., Oliva, A., Castelhano, M. S., & Henderson, J. (2006). Contextual guidance of eye movements and attention in real-world scenes: The role of global features in object search. Psychological Review, 113 (4), 766–786.

VanRullen, R., & Thorpe, S. J. (2001). The time course of visual processing: From early perception to decision-making. Journal of Cognitive Neuroscience, 13 (4), 454–461.

Võ, M. L.-H., & Wolfe, J. M. (2013). Differential Electrophysiological Signatures of Semantic and Syntactic Scene Processing. Psychological Science, 24 (9), 1816–1823.

Walther, D. B., & Shen, D. (2014). Nonaccidental Properties Underlie Human Categorization of Complex Natural Scenes. Psychological Science, 25 (4), 851–860.

Watson, D. M., Andrews, T. J., & Hartley, T. (2017). A data driven approach to understanding the organization of high-level visual cortex. Scientific Reports, 7 (1), 3596.

Watson, D. M., Hartley, T., & Andrews, T. J. (2014). Patterns of response to visual scenes are linked to the low-level properties of the image. NeuroImage, 99, 402–410.

Xiao, J., Ehinger, K. A., Hays, J., Torralba, A., & Oliva, A. (2014). SUN Database: Exploring a Large Collection of Scene Categories. International Journal of Computer Vision, 1–20.

Zheng, C. Y., Pereira, F., Baker, C. I., & Hebart, M. N. (2019). Revealing interpretable object representations from human behavior. arXiv:1901.02915 [cs, q-bio, stat]. Retrieved from http://arxiv.org/abs/1901.02915

Zhou, B., Lapedriza, A., Khosla, A., Oliva, A., & Torralba, A. (2017). Places: A 10 million Image Database for Scene Recognition. IEEE Transactions on Pattern Analysis and Machine Intelligence, 1–1.

